# Robust MR-AIV: A Systematic Study of Robustness Improvement and Sensitivity Analysis of MR-AIV

**DOI:** 10.64898/2026.04.14.718498

**Authors:** Mohammad Vaezi, Juan Diego Toscano, Yisen Guo, Ryszard Stefan Gomolka, George Em. Karniadakis, Douglas H. Kelley, Kimberly A. S. Boster

## Abstract

Cerebrospinal and interstitial fluid transport play a central role in brain metabolic waste clearance, yet non-invasive quantification of deep-brain flow dynamics remains challenging. Magnetic Resonance Artificial Intelligence Velocimetry (MR-AIV) is a physics-informed neural network framework that infers three-dimensional velocity, pressure, and permeability fields from dynamic contrastenhanced MRI by embedding porous-media flow physics into the learning process. Here, we present a methodological refinement and systematic evaluation of MR-AIV. We introduce a universal, anatomically informed, region-of-interest–based permeability initialization that improves anatomical alignment and physical consistency across subjects. We quantify the sensitivity of inferred fields to key modelling choices, including initialization strategies, permeability bounds, diffusivity assumptions, signal–concentration relationships, and measurement noise. Across these conditions, MR-AIV yields stable velocity and permeability estimates with preserved spatial structure. Together, these results establish practical guidelines and identify stable operating regimes for reliable deployment of MR-AIV. By improving robustness and reproducibility, this work strengthens MR-AIV as a minimally invasive approach for mapping brain-wide porous fluid transport and supports its application to studies of neurological health and disease.

## Introduction

### Brain Fluid Transport: Importance and Challenges in Its Estimation

Cerebrospinal fluid (CSF) and interstitial fluid (ISF) transport play a fundamental role in maintaining brain homeostasis with the removal of metabolic waste products via clearance pathways often referred to as the glymphatic system Xie et al. [2013], Rasmussen et al. [2018], Plog and Nedergaard [2018]. This transport occurs through a complex interplay of perivascular spaces (PVSs) and ISF spaces, where both advection (bulk flow) and diffusion likely contribute to solute movement Kelley and Thomas [2023], Ray et al. [2019], Guo et al. [2025]. The dysfunction of this system and the subsequent accumulation of large protein aggregates have been linked to neurological diseases, including Alzheimer’s disease and other neurodegenerative disorders Xie et al. [2013], Rasmussen et al. [2018], Plog and Nedergaard [2018], Mestre et al. [2020], Kelley and Thomas [2023].

The balance between fluid transport manifested by ISF influx and efflux remains an active area of investigation, particularly in deep brain regions where direct measurements are scarce. Impaired brain fluid transport has been associated with ageing Plog and Nedergaard [2018], Mestre et al. [2020], stroke Mestre et al. [2020], cerebral amyloid angiopathy Chen et al. [2022], Koundal et al. [2024], and hypertension Mestre et al. [2018]. Thus, understanding brain-wide transport mechanisms is critical not only for basic physiology but also for identifying early biomarkers of dysfunction and guiding therapeutic interventions. Eventually, fully noninvasive methods capable of quantifying brain fluid flow could enable patient-specific assessment of disease progression and treatment response.

Despite its importance, direct in vivo measurement of velocity, pressure, and permeability within brain tissue remains infeasible. Optical methods such as particle tracking velocimetry (PTV) provide high-fidelity velocity measurements Mestre et al. [2018], Min Rivas et al. [2020], Raghunandan et al. [2021], especially when enhanced with artificial intelligence velocimetry (AIV) Boster et al. [2023], Toscano et al. [2024], but they are restricted to small superficial regions accessible only via invasive cranial windows and do not provide brain-wide coverage. Front-tracking approaches applied to tracer (dye) imaging Mestre et al. [2020], Plog et al. [2018], Munk et al. [2019] allow for broader spatial estimation but neglect diffusion, are sensitive to thresholding choices, and generally lack mass conservation constraints.

Various MR approaches can provide insight into brain fluid mobility, but MR cannot directly measure velocities in the deep brain. Dynamic contrast-enhanced magnetic resonance imaging (DCE-MRI) enables whole-brain tracer imaging but requires invasive tracer infusion and is more applicable for measuring relative changes in contrast levels rather than velocity or pressure Valnes et al. [2020], Ray et al. [2021], Zapf et al. [2022]. Fully non-invasive approaches include using phase-contrast (PC) MRI to measure blood velocities in the large brain vessels or CSF at the level of the cerebral aqueduct. However, measuring the orders-of-magnitude-slower flows that occur throughout the brain, which are the primary focus of this work, is less reliable. Arterial spin labelling (ASL), with predominant application for cerebral perfusion imaging, can potentially quantify slow perfusion without contrast agents Hirschler et al. [2018], Pires Monteiro et al. [2025], and it has also been used to quantify the blood-to-CSF water transport in mice Lee et al. [2022]. Diffusion-weighted imaging (DWI) has also been used to estimate CSF and ISF mobility Wright et al. [2024]), and spectral DWI was shown to separate slow and fast water motion compartments in murine brain Gomolka et al. [2026]. However, diffusive motion does not accurately measure advection, and the vast majority of non-invasive MR methods, including ASL and DWI, inherently suffer from low spatial resolution, highlighting that DCE-MRI remains a predominant state-of-the-art in preclinical glymphatic research.

Computational approaches also face limitations. Optimal mass transport (OMT)–based methods estimate flow velocities from DCE-MRI measurements by incorporating diffusion and noise through regularization but rely on heuristic diffusivity-like parameters that lack clear physical interpretation and cannot directly infer permeability or pressure Ratner et al. [2017], Frangi et al. [2018], Koundal et al. [2020], Chen et al. [2022], Koundal et al. [2024]. Adjoint-based parameter estimation techniques depend on prescribed governing equations and do not typically reconstruct spatially heterogeneous permeability fields Vinje et al. [2023]. More broadly, many existing techniques either omit key physical constraints or cannot recover quantities such as permeability and pressure, which are central to porous transport governed by Darcy-type laws Holter et al. [2017], Bedussi et al. [2018], Basser [1992].

Brain fluid transport is inherently multiscale. Fast flows occur in large PVSs surrounding major arteries, while slow interstitial transport occurs within the porous interstitial space Ray et al. [2019], Guo et al. [2025]. The relative contributions of advection and diffusion vary spatially and temporally and can be characterized by local Péclet numbers Kelley and Thomas [2023], Guo et al. [2025]. Transport processes span several orders of magnitude in velocity and permeability, complicating both experimental measurement and computational modelling. Moreover, deep brain regions remain particularly inaccessible to direct probing, leaving critical aspects of intracranial transport poorly characterized. These challenges motivate the development of robust, physics-informed inverse frameworks capable of inferring physically meaningful transport quantities like velocity, pressure, and permeability, from indirect imaging data and tracer concentration Raissi et al. [2019, 2020].

### MR-AIV and the Need for Robustness

To address these challenges, Magnetic Resonance Artificial Intelligence Velocimetry (MR-AIV) was recently developed Toscano et al. [2025a]. It is a physics-informed machine learning framework designed to infer three-dimensional velocity, pressure, and permeability fields from DCE-MRI–derived tracer dynamics. Building on the principles of Artificial Intelligence Velocimetry (AIV) Boster et al. [2023], Toscano et al. [2024], Cai et al. [2021], Toscano et al. [2025b], MR-AIV integrates porous-media flow physics with observational data to solve the underlying advection–diffusion inverse problem Raissi et al. [2019, 2020]. By embedding Darcy’s law and the continuity equation within the neural network architecture and employing advanced variational frameworks, such as residual-based attention weights Anagnostopoulos et al. [2024a], Toscano et al. [2026], MR-AIV can reconstruct physically consistent flow fields from sparse and noisy concentration measurements.

While the feasibility of mapping deep-brain fluid velocities using physics-informed deep learning has been demonstrated Toscano et al. [2025a], the reliable application of such frameworks requires a rigorous understanding of their sensitivity to modelling assumptions, data quality, and user-defined parameters. Inverse problems in fluid mechanics are often ill-posed or sensitive to initialization Psaros et al. [2023]. Therefore, a comprehensive robustness analysis is essential before MR-AIV results can be confidently interpreted in biological or clinical contexts.

### Knowledge Gap: Sensitivity Analysis of MR-AIV

The sensitivity of MR-AIV is rooted in the ill-posed nature of the inverse problem and the intrinsic coupling between permeability and velocity fields through Darcy’s law. Because the governing equations link permeability gradients to velocity and pressure in a nontrivial and spatially heterogeneous manner, formal analytical sensitivity bounds are not tractable. Consequently, we adopt an empirical approach to quantify sensitivity across a range of practically relevant uncertainties. This perspective allows us to identify stable regimes and failure modes.

Although MR-AIV provides a powerful framework for inferring fluid dynamics quantities, several potential sources of uncertainty must be characterized:

- **Epistemic uncertainty:** MR-AIV relies on initial guesses for velocity and permeability fields and the learning range of permeability to guide the optimization process. Given the non-convex nature of the loss landscape in physics-informed neural networks Toscano et al. [2026], Anagnostopoulos et al. [2024b], the final prediction may depend on these initializations.
- **Model-form uncertainty:** Physical coefficients such as diffusivity are typically prescribed rather than learned. Although diffusivity is often assumed constant, substantial differences exist between open fluid spaces (e.g., PVSs) and porous interstitial space, and smaller spatial variations may also occur within tissue Kelley and Thomas [2023], Valnes et al. [2020].
- **Aleatoric uncertainty:** DCE-MRI data are affected by noise, limited temporal resolution, and assumptions regarding the conversion of signal enhancement ratio (SER) to tracer concentration Ratner et al. [2017], Stanton et al. [2021]. These factors introduce aleatoric uncertainty that may propagate to the predicted transport parameters Toscano et al. [2024], Psaros et al. [2023].

### Objectives and Scope of This Study

The objective of this work is to present a comprehensive sensitivity analysis and to enhance the robustness of the MR-AIV framework. We systematically assess the sensitivity of the framework to:

- **Initial permeability guesses:** We evaluate how different initializations affect the converged solution and introduce a standardized initialization strategy to improve robustness.
- **Initial velocity guesses:** We quantify the dependence of the predicted velocity fields on the initial estimates provided by front-tracking or uniform initial guesses.
- **Prescribed permeability learning ranges:** We analyse the impact of constraining the permeability search space on the final prediction.
- **Assumed SER–concentration relationships:** We test the robustness of the model to variations in the signalto-concentration mapping.
- **Spatial heterogeneity of diffusivity:** We explore how assumptions about homogeneous versus heterogeneous diffusivity affect the predicted pressure and velocity fields.
- **Measurement noise:** We assessed how measurement noise propagates to the predicted quantities.

By rigorously quantifying these sensitivities, we aim to establish best-practice guidelines for applying MR-AIV and to strengthen confidence in its reliability as a minimally invasive tool for studying brain-wide fluid transport.

## Methods

### Real and Synthetic Data

The experimental data for this study are comprised of time-resolved DCE-MRI time series of tracer transport throughout the wild-type mouse brain. Following the injection of tracer to the cisterna magna, three-dimensional T1-weighted gradient-spoiled gradient-echo (FISP) images were acquired with one-minute temporal resolution, enabling indirect observation of spatiotemporal tracer dynamics. The datasets analyzed in this study were previously reported, along with descriptions of acquisition parameters, tracer administration protocols, and animal preparation procedures Toscano et al. [2025a]. For each voxel, the primary observable quantity is the signal enhancement ratio *SER* = (*S*_*t*_ − *S*_0_)*/S*_0_ × 100 where *S*_*t*_ is the measured signal intensity at time *t* and *S*_0_ is the baseline voxel signal before tracer injection. The SER provides a normalized measure of signal change relative to baseline and serves as the input to the inverse framework. Because the governing transport model is expressed in terms of tracer concentration, the SER is converted to concentration using a prescribed SER–concentration relationship. In the baseline configuration, a linear mapping is used. While alternative nonlinear mappings are introduced below to evaluate robustness to signal-model uncertainty, it is worth highlighting that the FISP acquisition was chosen due to its ability to acquire higher signal amplitudes than other gradient-echo methods and with almost unchanged T1 magnetization across phase-encoding Scheffler and Lehnhardt [2003] during imaging. However, considering complexity of image contrast mechanism in FISP, defining a direct relation between the signal amplitude and gadolinium contrast concentration is not straightforward and is beyond the scope of the current study.

In DCE-MRI experiments, the true permeability, intracranial pressure, and velocity are unknown, making direct quantitative validation of reconstruction methods challenging or impossible. To rigorously evaluate the performance of the proposed framework, we therefore employ synthetic datasets. Specifically, computational fluid dynamics (CFD) simulations are used to generate synthetic data from permeability fields (chosen a priori) and appropriate boundary conditions, with flow governed by Darcy’s law and the continuity equation, and tracer motion governed by the advection-diffusion equation. The simulation was performed in a three-dimensional brain geometry reconstructed from a T2-weighted structural MRI using the finite element solver COMSOL Multiphysics 6.3. Darcy’s law was solved to obtain a steady velocity and pressure field under prescribed pressure boundary conditions (constant pressure at an inlet near the cisterna magna and zero pressure at designated outlet regions). The time-dependent advection–diffusion equation was then solved to generate transient tracer concentration fields, with a constant inlet concentration applied for the first 5 minutes and zero concentration thereafter. Thus, the permeability, velocity, and pressure were steady in time, while the tracer concentration evolved dynamically. The full simulation pipeline and synthetic data generation procedures are described in detail in Toscano et al. [2025a].

### Governing Physical Model

Tracer transport in brain tissue was modelled using the advection-diffusion equation. We considered two formulations depending on the diffusivity assumption.

#### Constant diffusivity

In the baseline formulation, diffusivity *D* was assumed uniform, yielding

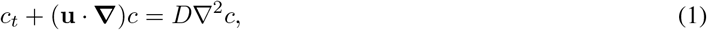

where *c*(**x**, *t*) denotes tracer concentration and **u**(**x**, *t*) denotes velocity. The solution is defined over the spatiotemporal domain Ω = Ω_x_ × Ω_*t*_ ⊂ ℝ^3+1^, with Ω_*t*_ = {*t* ℝ | *t*_0_ ≤ *t* ≤ *t*_*f*_}. Here, *t*_*f*_ = 90 minutes corresponds to the final acquisition time, and *t*_0_ = 10 minutes is selected to exclude the initial injection phase, since the source term in the transport equation is unknown during that interval.

#### Spatially varying diffusivity

In the second formulation, diffusivity was allowed to vary spatially, *D* = *D*(**x**), and the governing equation became

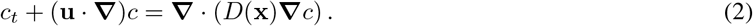

Compared to Eq. 1, this representation naturally introduces an additional term **∇***D* · **∇***c* when expanded, accounting for spatial heterogeneity in diffusion. In both formulations, *D* remained unchanged over time, and the flow was assumed incompressible, so that conservation of mass implies

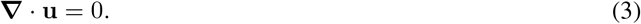

We adopted a porous-medium description of transport and imposed Darcy’s law:

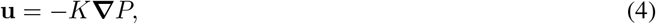

where *P* is pressure and *K* denotes hydraulic permeability, related to intrinsic permeability *κ* and dynamic viscosity *µ* through *K* = *κ/µ*. The velocity field was assumed steady over the anatomical domain Ω_x_ = {**x** ∈ ℝ^3^ **x** ∈ *G*_*m*_}, where *G*_*m*_ defines the brain geometry of mouse *m*. Substituting Eq. 4 into the transport and mass conservation equations yields, for the uniform-diffusivity case:

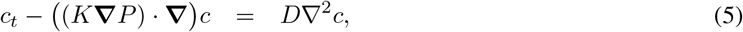

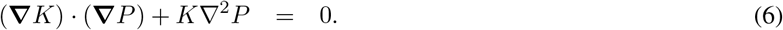

For the spatially varying diffusivity formulation, Eq. 5 is replaced by

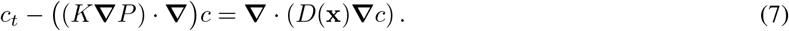

Because *c* is observed and *D* (or *D*(**x**)) is prescribed, this reformulation reduces the inverse problem to inferring two spatial fields, *K* and *P*, instead of three velocity components, thereby substantially mitigating the ill-posed problem. The permeability field *K* serves as a learned spatial map that captures sharp transitions between low- and high-velocity regions spanning multiple orders of magnitude. Once *K* and *P* are predicted, the velocity field **u** is estimated via Eq. 4.

All governing equations were non-dimensionalised prior to training to improve numerical stability, as physical parameters span several orders of magnitude (e.g., *κ* ∼ 10^−10^ mm^2^). Non-dimensionalisation prevents loss of gradient information due to single-precision arithmetic and is standard practice in physics-informed machine learning frameworks Toscano et al. [2025c], Wang et al. [2023]. Non-dimensionalization was performed using characteristic scales defined by a representative domain length *L*_char_, a characteristic velocity *U*_char_, the associated advective time scale *T*_char_ = *L*_char_*/U*_char_, a reference diffusivity *D*_char_, and a reference permeability *κ*_char_. Pressure was scaled by *P*_char_ = *µU*_char_*L*_char_*/κ*_char_, and concentration was normalized by a unit reference value. The complete non-dimensional equations are provided in Toscano et al. [2025a].

### MR-AIV Framework

MR-AIV is a physics-informed neural network (PINN) framework designed to infer velocity, pressure, and permeability fields from concentration data, as sketched in Fig. 1. Two neural networks are designed to separate noise from mean-concentration data, and two other networks approximate pressure and permeability. In all networks, spatial coordinates **x** and time *t* serve as inputs. The permeability field is parameterized as a spatially varying but time-independent output. When using a spatially varying prescribed diffusivity, we define another network for this variable. Training is performed by minimizing a total loss function consisting of

1. a data mismatch term enforcing agreement between predicted and measured concentration,
2. a residual term enforcing the advection–diffusion equation and Darcy’s law (either Eq. 5 or Eq. 7), and
3. a divergence penalty enforcing conservation of fluid mass (Eq. 6).

**Figure 1:**
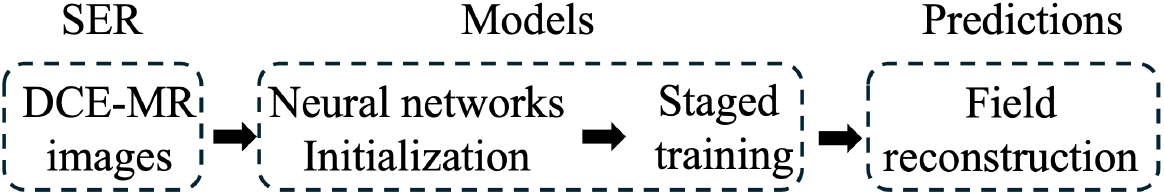
MR-AIV framework. MR-AIV is a framework designed to solve the inverse problem of predicting velocities from concentrations. The SER data derived from DCE-MR images are used to train physics-informed models in two main steps: first, network initialization, and second, staged training. The trained models are then used to predict pressure, permeability, and velocity. Toscano et al. [2025a].

Automatic differentiation is used to compute spatial and temporal derivatives required for the governing equations. All residual terms are evaluated at collocation points distributed throughout the spatiotemporal domain. Pressure and permeability are predicted simultaneously through joint optimization. Permeability is constrained to lie within a prescribed range during training. Optimization is continued until the total loss function converges and the predicted fields stabilize. No direct supervision of permeability or pressure estimation is used; these quantities are predicted solely through consistency with measured concentration dynamics and governing physics Toscano et al. [2025a].

### Sensitivity Analysis Design

To evaluate robustness, we systematically perturbed model inputs, assumptions, and training configurations. Each perturbation was applied independently while holding all other settings fixed. To quantitatively compare the predicted fields obtained under different modelling assumptions, we computed two complementary metrics: the Wasserstein distance and the gradient-based structural similarity index (G-SSIM). The Wasserstein distance quantifies differences between the distributions of predicted speeds, while G-SSIM evaluates the similarity of spatial structures in the velocity fields. Smaller Wasserstein distances and larger G-SSIM values indicate stronger agreement between predictions. The pairwise comparisons for all tested configurations are summarized in Table 1.

**Table 1:** Pairwise comparison of predicted speed fields obtained under different modeling choices. For each comparison matrix, the Wasserstein distance (upper triangular entries) quantifies differences between the distributions of predicted speed, while the G-SSIM (lower triangular entries) measures similarity in the spatial organization of the predicted speed fields. Smaller Wasserstein distances and larger G-SSIM values indicate stronger agreement between predictions. (a) Effect of velocity initialization strategies (constant, FT1, FT2). (b) Effect of permeability initialization (binary, 10ROI, 92ROI, and perturbed 92ROI). (c) Effect of the SER–concentration model formulation (linear vs nonlinear). (d) Effect of the permeability training range (4-decade vs 5-decade range). (e) Effect of the diffusivity prescription (D1, D2, D1(x), and D2(x)).

| **(a) Velocity initialization** |  |  |  |  |
| --- | --- | --- | --- | --- |
|  | Prediction constant | Prediction FT1 | Prediction FT2 |  |
| Prediction constant | – | 0.097 | 0.094 |  |
| Prediction FT1 | 0.905 | – | 0.009 |  |
| Prediction FT2 | 0.917 | 0.910 | – |  |
| **(b) Permeability initialization** |  |  |  |  |
|  | Prediction binary | Prediction 10ROI | Prediction 92ROI | Prediction perturbed 92ROI |
| Prediction binary | – | 0.327 | 0.224 | 0.190 |
| Prediction 10ROI | 0.918 | – | 0.104 | 0.138 |
| Prediction 92ROI | 0.906 | 0.909 | – | 0.035 |
| Prediction perturbed 92ROI | 0.906 | 0.910 | 0.925 | – |
| **(c) Permeability training range** |  |  |  |  |
|  | Prediction 4-decades | Prediction 5-decades |  |  |
| Prediction 4-decades | – | 0.128 |  |  |
| Prediction 5-decades | 0.910 | – |  |  |
| **(d) SER-concentration model** |  |  |  |  |
|  | Prediction linear concentration | Prediction nonlinear concentration |  |  |
| Prediction linear concentration | – | 0.087 |  |  |
| Prediction nonlinear concentration | 0.912 | – |  |  |
| **(e) Diffusivity prescription** |  |  |  |  |
|  | Prediction D1 | Prediction D2 | Prediction D1(x) | Prediction D2(x) |
| Prediction D1 | – | 0.217 | 0.389 | 0.351 |
| Prediction D2 | 0.911 | – | 0.173 | 0.134 |
| Prediction D1(x) | 0.902 | 0.913 | – | 0.039 |
| Prediction D2(x) | 0.900 | 0.913 | 0.921 | – |

#### Initial permeability guess

The MR-AIV framework requires an initial guess for the permeability field. Though the permeability evolves through the course of training the PINN model, we wondered how much the final, predicted permeability might be affected by the initial guess. Four strategies were tested for constructing an initial permeability guess. In the first strategy, described previously Toscano et al. [2025a], regions where contrast became enhanced within 10 minutes of injection were assigned an initial intrinsic permeability *κ* = 10^−6^ mm^2^ and other regions were assigned *κ* = 10^−10^ mm^2^, yielding a binary map of the permeability field of the brain. After training, the final predicted permeability field was projected onto a standard brain atlas (electronic supplementary material S1). As an initial step, we computed the median permeability in each of 10 regions of interest (ROIs) (electronic supplementary material S1), consistent with the analysis reported in **?**: Hippocampus (HIP), Caudate nucleus (CP), Thalamus (TH), Midbrain (mesencephalon MB), Superior sagittal sinus (SSS), Perivascular spaces (PVS) of the Circle of Willis (PVS–CoW), Perivascular spaces of the Middle Cerebral Artery (PVS–MCA), Perivascular spaces of the Anterior Cerebral Artery (PVS–ACA), Perivascular spaces of the Basilar Artery (PVS–basilar), and Subarachnoid space of the olfactory region (SAS–olfactory). This procedure was repeated for four additional wild-type mice, altogether yielding five subject-specific atlas-region-based permeability maps, each containing permeability estimates for the same 10 ROIs. For each mouse, the median permeability of the remaining brain tissue was treated as the remainder region. The median (instead of mean) was calculated to account for possible spatial registration inaccuracies, especially in regions of small voxel count. This resulted in five permeability maps, each consisting of 11 regions (10 ROIs plus the remainder of the brain). We then computed the mean of the regional medians across the five mice to construct a consolidated map, referred to as the *universal 10ROI permeability map*. This map was subsequently used as an alternative initialization for the permeability field — the second strategy. The same procedure was repeated using 92 ROIs from the atlas, leading to the construction of a *universal 92ROI permeability map* (electronic supplementary material S1). Using the 92ROI map as the initial permeability guess was the third strategy. For the fourth strategy, we generated a perturbed version of the 92ROI map by applying multiplicative Gaussian noise in linear space according to

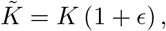

where *ϵ* ∼ 𝒩 (0, *α*^2^) with *α* = 0.15. This corresponds to 15% zero-mean relative variability at each spatial location, such that the mean permeability is preserved (electronic supplementary material S1). These different initializations were deliberately designed to exhibit substantially different spatial permeability distributions, enabling assessment of convergence behaviour and robustness of the proposed framework.

#### Initial velocity guess

The MR-AIV framework also requires an initial guess for the velocity field, which we produced from front tracking. In front tracking, fronts, or regions of constant concentration, are tracked over time and space Toscano et al. [2025a]. Front tracking neglects diffusion and thus has low accuracy in regions where diffusion plays a substantial role in transport. We wondered how much the final, predicted velocity might be affected by the initial guess. Three strategies were tested for constructing an initial velocity guess (electronic supplementary material S3):

- Two front-tracking–based estimates, denoted FT1 and FT2, were obtained from DCE-MRI data by applying different SER threshold intervals: [25%, 75%] for FT1 and [15%, 95%] for FT2.
- In lieu of front tracking, a spatially uniform velocity field of constant magnitude (*u, v, w* = 0.1 mm/min) was used as the initial guess.

#### Permeability learning range

The MR-AIV framework requires lower and upper bounds on the permeability during training. Previously, brain permeability was allowed to vary over four orders of magnitude, from 10^−10^ to 10^−6^ mm^2^ Toscano et al. [2025a]. In this study, we allowed the permeability to vary over five orders of magnitude, from 10^−11^ to 10^−6^ mm^2^, to assess the influence of domain constraints on the solution.

#### SER–concentration mapping

Both linear and nonlinear (cubic polynomial derived from Ratner et al. [2017]) SER–concentration relationships were tested to assess sensitivity to signal-model uncertainty.

#### Diffusivity assumptions

Diffusivity was treated as prescribed and fixed during training. The original implementation of MR-AIV used a constant diffusivity, but the effective diffusivity in a porous medium *D*_eff_ is known to vary as *D*_eff_ = *D*_free_*/λ*^2^, where *D*_free_ and *λ* are the free diffusivity and tortuosity, so we expect that the diffusivity of Gd-DPTA tracer in the brain varies between *D*_free_ = 3.8 × 10^−4^ mm^2^/s (the free diffusion coefficient for Gd-DPTA, which has a similar molecular weight as gadobutrol) Valnes et al. [2020] and *D*_eff_ = 1.48 × 10^−4^ mm^2^/s (assuming *λ* = 1.6). We investigated how the prescribed diffusivity affects the predicted results. We first considered two spatially uniform diffusivities, *D*_1_ = 2.4 × 10^−4^ mm^2^/s (the same value used in Toscano et al. [2025a]) and *D*_2_ = 1.847 × 10^−4^ mm^2^/s, to assess sensitivity to the prescribed mean diffusivity. To investigate the impact of spatial heterogeneity, we constructed two spatially varying diffusivity fields with the same mean values *D*_2_. The first spatially varying diffusivity field, denoted *D*_1_(**x**), was constructed based on the predicted speed obtained using the spatially uniform diffusivity *D*_1_. Specifically, the speed ||**u**|| was first computed and log_10_(||**u**||) was rescaled to linear range [10^−7^, 6.21 × 10^−4^] mm^2^/s. The resulting diffusivity field was continuous (not binary), and no spatial smoothing was applied. Regions with larger speed were therefore assigned larger diffusivity values through this mapping, while regions with lower speed were assigned smaller diffusivity values. The final field was rescaled to ensure that its spatial mean was *D*_2_ = 1.847 × 10^−4^ mm^2^/s. The second spatially varying diffusivity field, *D*_2_(**x**), was constructed based on early tracer arrival patterns. Specifically, a map from early tracer arrival Toscano et al. [2025a] was normalized to the range [0, 1] then linearly rescaled to [1.18 × 10^−4^, 1.18 × 10^−3^] mm^2^/s and the mean of final diffusivity field matched mean(*D*_2_(*x*)) = *D*_2_ = 1.847 × 10^−4^ mm^2^/s (electronic supplementary material S10).

#### Measurement noise

To assess sensitivity to measurement noise, synthetic concentration data were contaminated with noise in four different cases: additive Gaussian noise with amplitudes equal to 10% and 50% of the mean concentration in the field, and sparse outliers with concentrations of five and two times the mean concentration, selected uniformly at random, comprising 5% and 2% of the domain, respectively. The model was retrained independently for each case.

## Results

### Universal ROI-based permeability initializations improve robustness and consistency of inferred fields

We first evaluated the sensitivity of the proposed framework to the choice of the initial permeability field. Four distinct initialization strategies were considered, as described above: a binary permeability map derived from early-time tracer arrival dynamics, a universal anatomically informed 10-region-of-interest (10ROI) map, a universal 92ROI map, and a perturbed version of the 92ROI map. These permeability distributions were constructed to differ substantially in order to rigorously assess the convergence and robustness of the model.

Figure 2 compares the initial and predicted permeability and speed distributions for all four initialization strategies. As shown in Fig. 2a, the initial permeability distributions differ markedly, with a bimodal distribution of binary initialization and unimodal distributions for anatomically informed ROI-based initialization maps.The 10ROI, 92ROI, and perturbed 92ROI initialization strategies yield similar predicted permeability distributions after training, though the binary initialization does not (Fig. 2b). Specifically, the binary initialization predicts a distribution with reduced bimodality.

**Figure 2:**
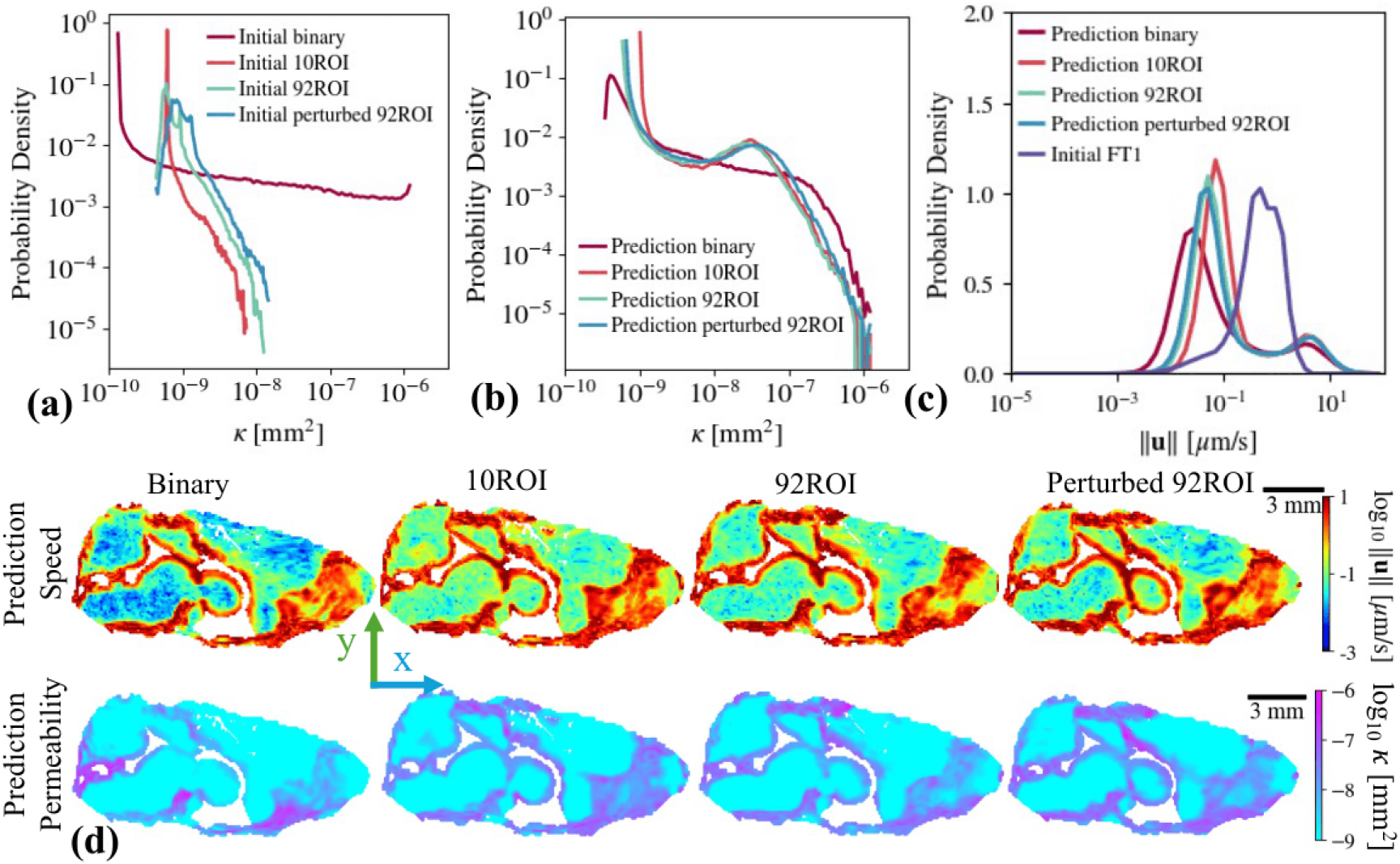
Universal ROI-based permeability initialization yields robust and consistent predictions. Four initialization strategies are compared: binary, 10ROI, 92ROI, and perturbed 92ROI. (a) Initial distributions of intrinsic permeability *κ*. (b) Predicted permeability distributions after training. ROI-based initializations converge to nearly identical solutions, while the binary initialization shows reduced bimodality. (c) Predicted distributions of speed ||*u*|| exhibit bimodal structure in all initialization strategies, with minor deviation for the binary initialization. (d) Midsagittal maps of predicted speed and permeability. Spatial patterns are consistent for ROI-based initializations. Together these results show that all of the ROI-based permeability guesses yield similar predictions, but the binary guess does not.

A similar trend is observed for the predicted speed distributions (Fig. 2c, Table 1b). Across all initialization strategies, the recovered speed fields exhibit a clear bimodal structure with fast and slow regions. The fast flow has a speed ∼3 µm/s in all initialization strategies, whereas the slow flow is slower when the binary initialization is used than when the ROI-based initializations are used. Notably, the 92ROI initialization and its perturbed counterpart produce nearly identical speed and permeability distributions, demonstrating the robustness of the model to spatial perturbations in the initial permeability guess. Spatial maps of the predicted speed and permeability fields (Fig. 2d, Table 1b, electronic supplementary material S4, S5) confirm these findings: the ROI-based initializations produce visually and quantitatively similar spatial patterns, whereas the binary initialization exhibits some deviation. Collectively, these results indicate that anatomically informed universal ROI-based initializations lead to similar and consistent permeability and speed estimates.

To further assess the quality of the recovered fields, we evaluated the anatomical and physical consistency of the predictions (Fig. 3). Anatomical consistency was assessed by comparing binarized and normalized spatial gradients of the predicted permeability field with binarized and normalized gradients of the anatomical reference T2-weighted MR images, which serve as a proxy for anatomical boundaries. To determine the appropriate binarization thresholds, we evaluated multiple threshold levels for both the permeability-based and MRI-based images across five wild-type mice. For each threshold combination, we computed the G-SSIM and the Intersection over Union (IoU) metric. Setting the threshold equal to the 97th percentile of the gradient for the MRI-based images and the 90th percentile for the permeability-based images produced the highest similarity scores. We observed regions of elevated permeability and velocity surrounding the ventricles in the predicted fields. To account for these high-permeability “buffer” regions, we dilated the binarized structural MRI image by 3 voxels to reflect that the spatial extent of elevated permeability exceeds the anatomical boundaries captured by the structural image. Comparisons of midsagittal, axial, and coronal views (Fig. 3a) show that permeability gradients obtained from the 10ROI and 92ROI initializations align more closely with structural boundaries than those obtained from the binary initialization. This observation is quantitatively supported by the G-SSIM and the IoU metrics (Fig. 3b–c). Across five wild-type mice, the universal 10ROI initialization yields significantly higher similarity scores than the binary initialization (*p* = 0.0047 for G-SSIM and *p* = 0.0142 for IoU, T-test), confirming improved anatomical consistency.

**Figure 3:**
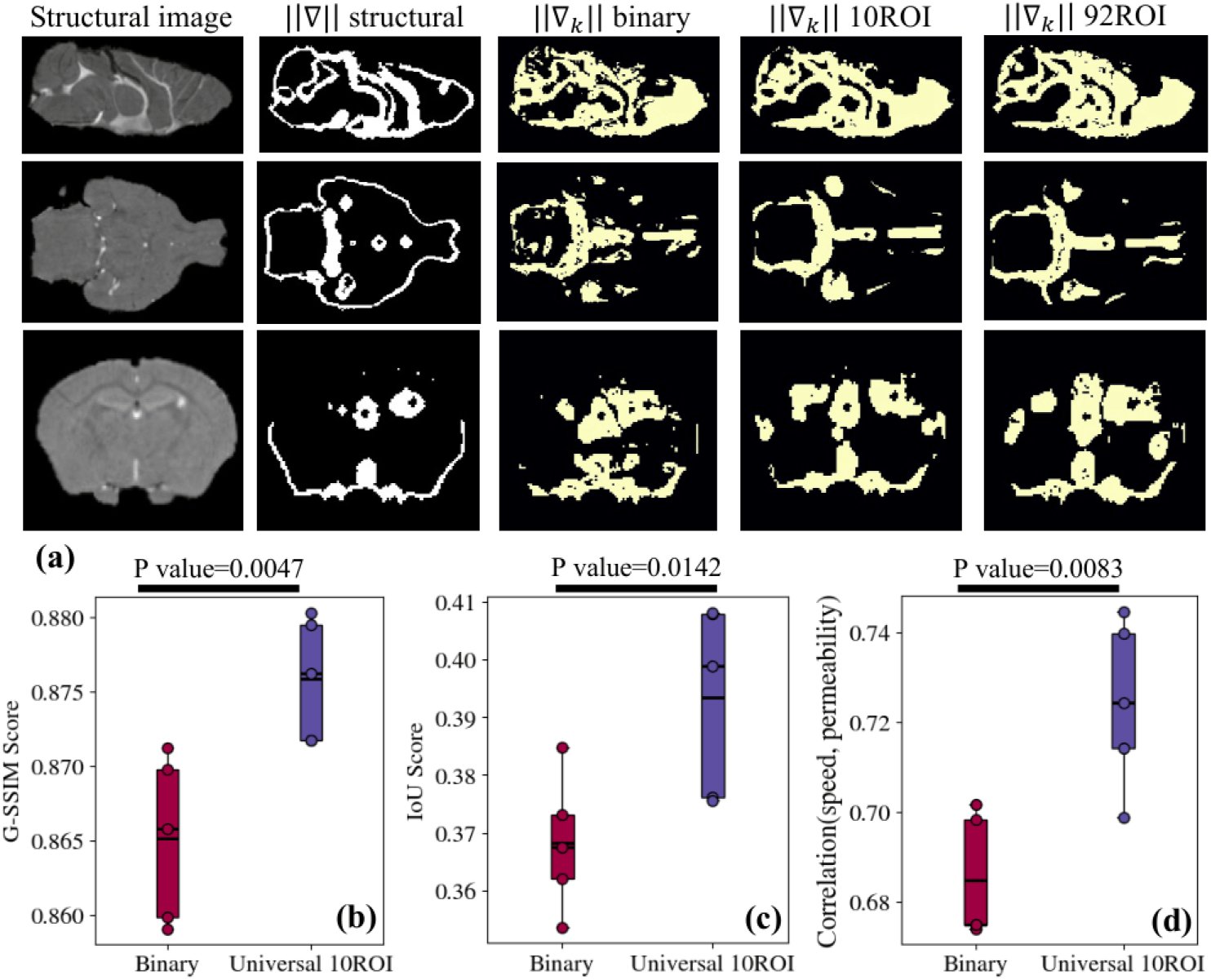
Universal ROI-based permeability initialization improves anatomical and physical consistency. (a) Comparison of structural MRI binarized and normalized gradients with gradients of predicted permeability for binary, 10ROI, and 92ROI initializations on three anatomical planes. The ROI-based initialization strategies show improved alignment with anatomical boundaries. (b) Gradient-based structural similarity index (G-SSIM) and (c) Intersection over Union (IoU) metrics across five mice confirm significantly higher anatomical consistency with the 10ROI initialization than with the binary initialization. (d) Speed–permeability correlation coefficients are higher, demonstrating improved physical consistency, with the 10ROI initialization than with the binary initialization. Together, these results support adoption of the universal 10ROI initialization.

Though gradients of intracranial pressure have rarely been measured, they are expected to be weak. Thus, for physical consistency with Eq. 4, we expect spatial variations of speed to correlate with those of permeability. Correlation coefficients, for the same five mice, are shown in Fig. 3d. The universal ROI-based initialization yields a significantly higher speed–permeability correlation than the binary initialization (*p* = 0.0083), indicating improved adherence to the governing physical model.

Taken together, these results demonstrate that universal ROI-based permeability initializations produce predictions that are significantly more consistent with both anatomical reference and expected physical flow relationships than binary initializations. Therefore, the universal 10ROI permeability map is adopted as the default permeability initialization strategy for the remainder of this study. We show the velocity and permeability distributions for the five mice in the electronic supplementary material S2. The results are similar to those reported by Toscano et al. Toscano et al. [2025a], but the inter-mouse variability is reduced, and the brain-wide speed distribution is more bimodal.

### Predicted results are robust to varying velocity initialization

We next evaluated the sensitivity of velocity predictions to the initial velocity guess. We considered three distinct velocity initialization strategies spanning a wide range of assumptions: two front-tracking–based velocity fields FT1 and FT2 derived using different thresholding criteria, and a spatially uniform velocity field with constant magnitude (*u, v, w* = 0.1 mm/min) throughout the whole brain domain.

Figure 4a shows the distributions of the initial magnitude and components of the velocity for the three initialization strategies, which differ substantially. Despite these pronounced differences, all three predicted velocity distributions converge to nearly identical forms after training (Fig. 4b). This convergence demonstrates that the optimization process consistently drives the solution toward a specific velocity field, largely independent of the initial guess.

**Figure 4:**
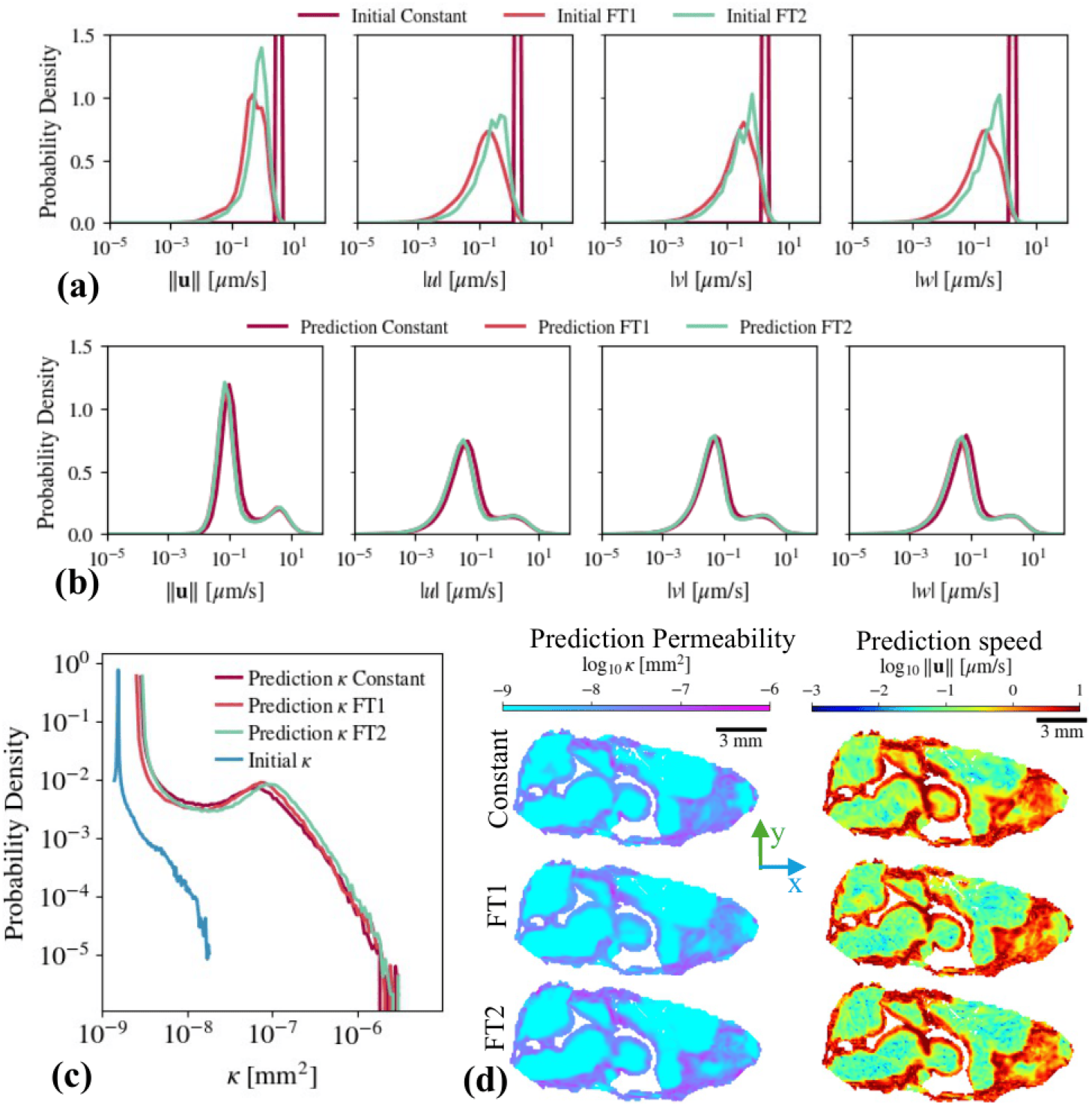
Velocity initialization does not affect predicted velocity or permeability. Three velocity initializations are compared: two front-tracking–based fields (FT1, FT2) and a spatially uniform field. (a) Initial distributions of speed and velocity components show substantial differences. (b) Predicted velocity distributions strongly overlap, indicating convergence to a consistent solution. (c) Predicted permeability distributions (using 10ROI permeability initialization) are nearly identical across velocity initializations. (d) Midsagittal slices of predicted permeability and speed show similar spatial patterns. These results demonstrate robustness of the method in velocity and permeability inference to velocity initialization.

We further examined whether velocity initialization influences permeability predictions (Fig. 4c). For all initialization strategies, the universal 10ROI permeability map was used as the initial permeability guess to isolate the effect of velocity initialization. The predicted permeability distributions corresponding to the three velocity initializations strongly overlap, indicating that prediction of permeability is insensitive to the choice of velocity initialization. This is particularly noteworthy given the coupling between velocity and permeability in the governing equations, and suggests that the joint inference procedure is well-constrained by the data and physical model.

Spatial comparisons reinforce these statistical findings. Midsagittal slices of the predicted permeability and speed magnitude (Fig. 4d, electronic supplementary material S6, S7) reveal highly similar spatial patterns across all initialization strategies. Both global distributional properties and local spatial features of the predicted fields are preserved, regardless of whether the initial velocity field is based on front tracking (FT1, FT2) or spatially uniform. These observations are supported quantitatively by the similarity metrics reported in Table 1a, which show small Wasserstein distances and high G-SSIM values between predictions obtained from different velocity initialization strategies.

Together, these results demonstrate that the proposed framework converges to consistent velocity and permeability solutions independent of the initial velocity guess. The method does not require a carefully tuned or highly accurate velocity initialization for reliable inference, highlighting its robustness and practical applicability.

### Predicted results are robust to expansion of the permeability learning range

We next investigated the sensitivity of the proposed framework to the prescribed permeability range. In the MR-AIV framework, lower and upper bounds on permeability must be imposed to constrain the optimization. Based on prior literature, permeability was initially restricted to vary over four decades, from 10^−10^ to 10^−6^ mm^2^. To evaluate robustness to this assumption, we expanded the allowable domain to five decades, from 10^−11^ to 10^−6^ mm^2^, and compared the resulting predicted velocity and permeability fields.

Figure 5 summarizes the comparison between the four- and five-decade ranges. Panel (a) shows the distributions of the initial and predicted magnitude and components of the velocity for each permeability range. The predicted velocity distributions corresponding to the four- and five-decade cases are consistent with each other but differ from the initial distribution. In both cases, the speed distribution exhibits a clear bimodal structure. While the high-speed peak is unchanged, and the low-speed peak shifts lower with a wider permeability range, without altering the overall spatial pattern of the recovered velocity field.

**Figure 5:**
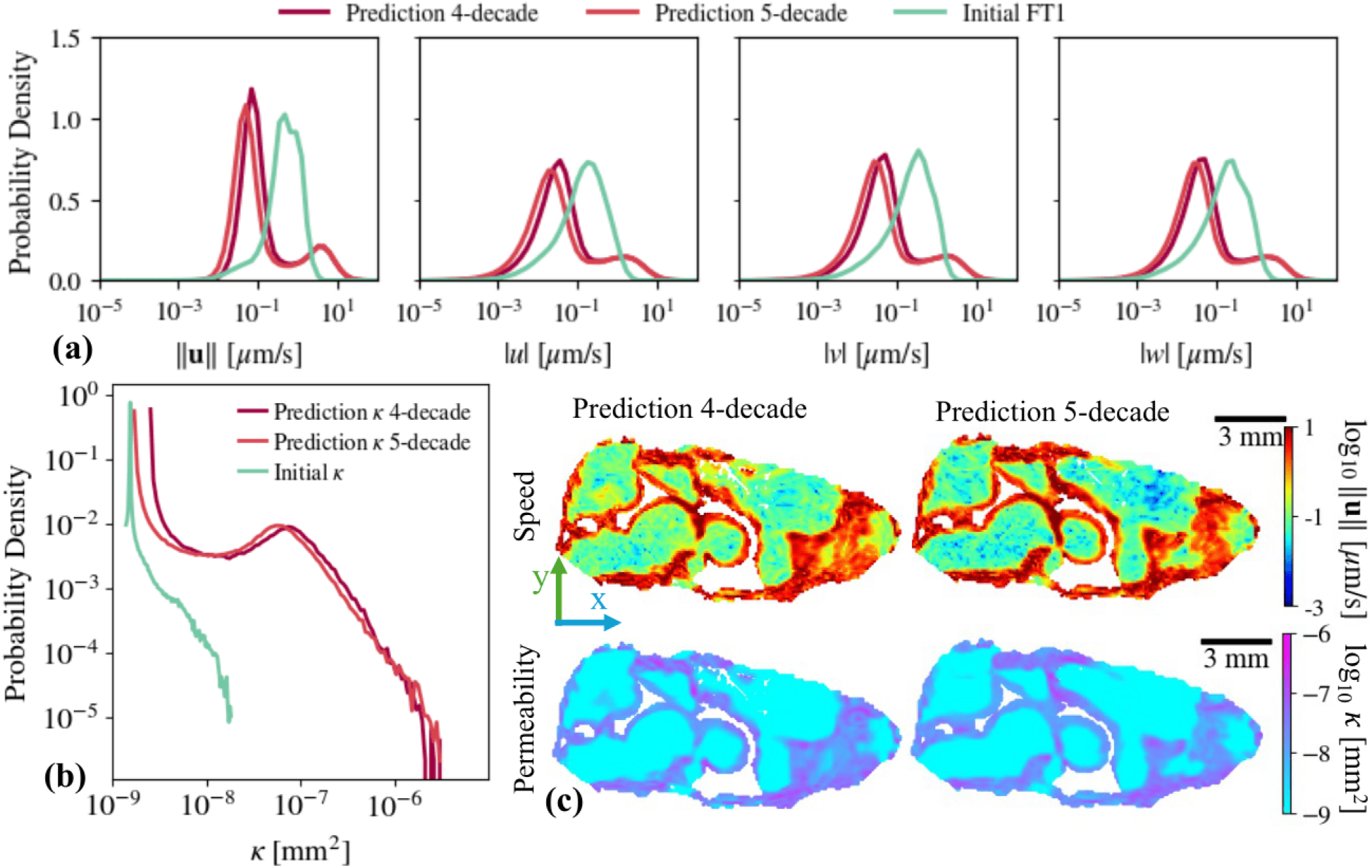
Expanding permeability bounds does not qualitatively change predicted fields. Model predictions are compared for permeability bounds spanning four and five decades. (a) Distributions of speed and velocity components show strong overlap, with only minor differences in the low-speed regime. (b) Permeability distributions are largely preserved, with a slight extension toward lower values in the five-decade case. (c) Midsagittal maps of predicted speed and permeability show similar spatial organization regardless of the permeability range. These results demonstrate that expanding the permeability range does not qualitatively alter predicted velocity or permeability fields.

Panel (b) compares the probability density functions of the initial permeability guess and the predicted permeability fields for the two ranges. Because the same initial permeability map is used in both cases, the comparison isolates the effect of the expanded learning domain. The predicted permeability distributions overlap closely, particularly in the high-permeability regime. A minor extension toward lower permeability values appears in the five-decade case, consistent with the shift observed in the low-speed region of the velocity distribution. Nevertheless, the overall structure, dynamic range, and bimodal characteristics of the permeability field remain preserved.

Spatial comparisons further confirm these observations. Panel (c) and electronic supplementary material S8 present slices of the predicted speed and permeability for both learning ranges. The spatial organization of high- and low-permeability regions, as well as the corresponding flow patterns, is similar between the four- and five-decade cases. The quantitative comparison in Table 1c also indicates strong agreement between the four- and five-decade permeability ranges, with small Wasserstein distance and high spatial similarity.

Together, these results demonstrate that the predicted velocity and permeability fields are robust to the permeability range. Expanding the range by one decade does not substantially alter either the statistical distributions or the spatial patterns of the recovered fields. Given this stability and consistency with literature-based expectations, the four-decade permeability range is adopted for the remainder of the study.

### Predicted results are robust to SER–concentration modelling assumptions

In DCE-MRI acquisitions, only the signal enhancement ratio (SER), expressed as percent enhancement relative to baseline, is directly measurable. Conversion from SER to tracer concentration, therefore, requires an assumed signal–concentration relationship. Because this conversion introduces modelling uncertainty, we evaluated its impact on the predicted velocity and permeability fields by comparing linear and nonlinear SER–concentration formulations.

In the linear case, SER values were mapped linearly to (dimensionless) concentrations ranging from 0 to 0.5, corresponding to the minimum and maximum SER observed in the dataset, respectively. In the nonlinear case, a cubic polynomial relationship, derived from the mapping in Ratner et al. [2017], was employed. Importantly, both mappings were applied uniformly across the dataset, and identical initial guesses for velocity and permeability were used in both cases to isolate the effect of the signal–concentration model.

Figure 6a compares the distributions of the magnitude and components of the velocity. In each case, the predicted velocity distributions nearly overlap and differ from the initial velocity distribution. This close agreement indicates that the predicted velocity field is largely insensitive to the assumed linearity or nonlinearity of the SER–concentration relationship, and that uncertainty in concentration estimation does not substantially propagate into velocity inference.

**Figure 6:**
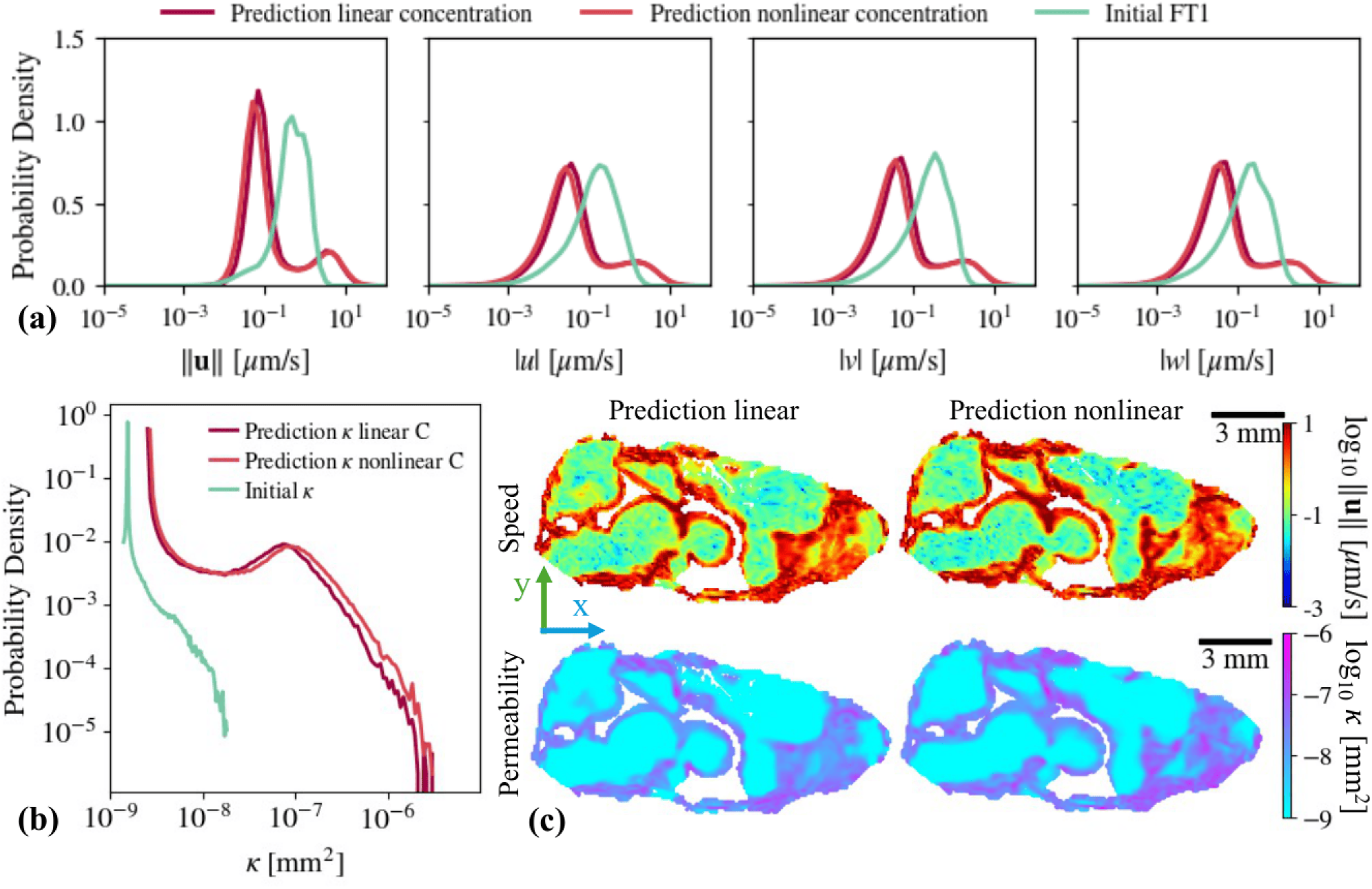
Linear and nonlinear SER–concentration mappings yield consistent predictions. Model predictions are compared using linear and nonlinear (cubic) SER–concentration mappings. (a) Distributions of speed and velocity components strongly overlap between the two mappings. (b) Permeability distributions are largely preserved, with a slight extension of the high-permeability tail under the nonlinear mapping. (c) Midsagittal maps of predicted speed and permeability show similar spatial organization with linear or nonlinear mapping. Moderate nonlinearity in SER–concentration conversion does not qualitatively affect predicted velocity or permeability fields.

Figure 6b presents the corresponding permeability distributions, which are also consistent between the linear and nonlinear cases. A modest extension of the high-permeability tail is observed with the nonlinear mapping. This shift is physically consistent with the choice of signal-concentration model: because the cubic relationship assigns slightly higher concentrations to high-SER voxels, it leads to correspondingly elevated permeability estimates in those regions. However, this difference remains systematic and small, without altering the overall structure or range of the distribution.

Spatial comparisons further confirm these findings. Midsagittal slices of the predicted speed and permeability fields (Fig. 6c and electronic supplementary material S9) show that spatial structures are consistent between the linear and nonlinear cases. High- and low-value regions are preserved, and no spatial artifacts or qualitative changes emerge due to the nonlinear mapping. Although the nonlinear case exhibits slightly more pronounced high-permeability regions, the overall spatial patterns remain essentially unchanged. Consistent with the qualitative comparisons in Fig. 6, the quantitative metrics in Table 1d show small Wasserstein distance and high G-SSIM between the linear and nonlinear SER–concentration mappings, indicating strong agreement between the resulting velocity fields.

Collectively, these results demonstrate that the proposed framework is robust to uncertainty in the SER–concentration relationship. Both linear and nonlinear mappings yield qualitatively and quantitatively similar velocity and permeability fields, indicating that moderate nonlinearity in signal–concentration conversion does not substantially influence predictions. Given this stability, a linear SER–concentration relationship is adopted for subsequent analyses.

### Predicted results are sensitive to mean diffusivity but robust to spatial heterogeneity

We next examined the influence of the assumed diffusivity on the predicted velocity and permeability fields. In the MR-AIV framework, diffusivity is prescribed and held fixed during training rather than learned. Therefore, it is important to assess sensitivity to the magnitude and spatial heterogeneity of the diffusivity.

Four diffusivity configurations were considered (Fig. 7). Two cases employed spatially uniform diffusivity values: *D*_1_ = 2.4 × 10^−4^ mm^2^*/*s and *D*_2_ = 1.847 × 10^−4^ mm^2^*/*s. To investigate spatial heterogeneity, two additional diffusivity fields were constructed while preserving the mean value 1.847 × 10^−4^ mm^2^*/*s. The first heterogeneous field, *D*_1_(**x**), was derived from a trained speed map. The second heterogeneous field, *D*_2_(**x**), was generated from early tracer arrival patterns.

**Figure 7:**
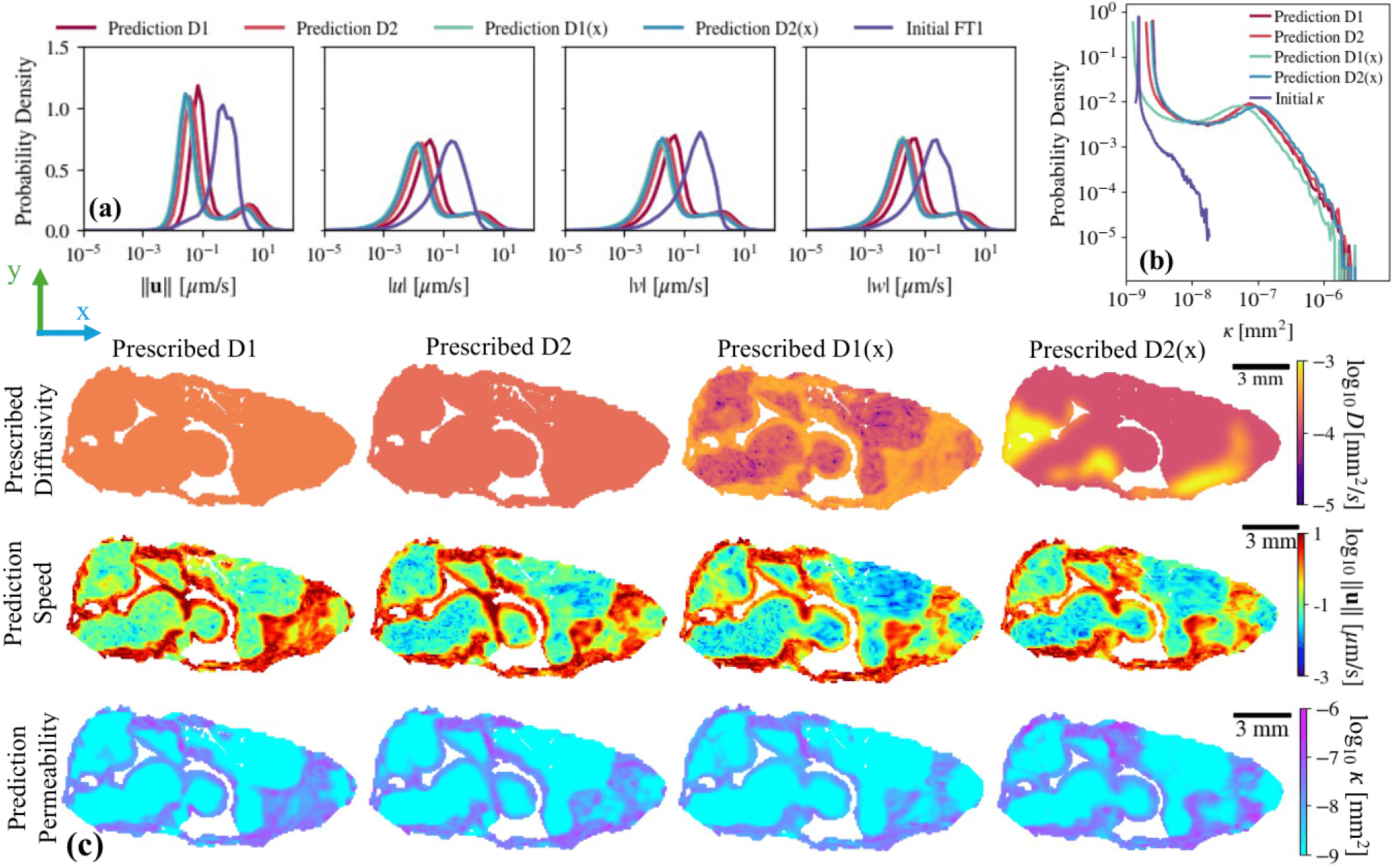
Predicted velocity and permeability are robust to spatial diffusivity variations. Four prescribed diffusivity configurations are compared: two spatially uniform cases with different mean values and two spatially heterogeneous cases with identical mean diffusivity. (a) Velocity distributions show modest sensitivity to changes in mean diffusivity but strong overlap among cases with the same mean. (b) Permeability distributions remain largely preserved across all configurations. (c) Midsagittal maps of diffusivity (top), predicted speed (middle), and predicted permeability (bottom) demonstrate consistent spatial organization of predicted fields even as diffusivity varies. These results indicate sensitivity to mean diffusivity but not to spatial variations of diffusivity.

Figure 7a shows the distributions of the initial and predicted magnitude and components of the velocity for each diffusivity assumption. Comparing the predictions obtained using *D*_1_ and *D*_2_ shows that decreasing the mean diffusivity lowers the low-speed peak of the distribution. However, the high-speed peak remains largely unchanged, and the overall bimodal structure of the speed distribution is preserved. Crucially, for the three cases sharing the same mean diffusivity (*D*_2_, *D*_1_(**x**), and *D*_2_(**x**)), the predicted velocity distributions nearly overlap across all components. This indicates that spatial heterogeneity in diffusivity does not substantially influence velocity predictions when the mean diffusivity is fixed.

The corresponding permeability distributions are shown in Fig. 7b. All cases exhibit similar structure and spread, with no emergence of new modes or qualitative changes. A slight shift toward lower permeability values is observed in the *D*_1_(**x**) case, consistent with the presence of localized lower-diffusivity regions in the prescribed map; because diffusivity was constructed to increase with speed, low-speed regions were assigned lower diffusivity, which weakens local diffusion and lowers the inferred permeability. Nevertheless, these differences are small and remain quantitative rather than structural. Overall, permeability prediction appears largely insensitive to spatial diffusivity variations at fixed mean diffusivity.

Spatial comparisons further support these conclusions. Figure 7c and electronic supplementary material S11 and S12 present slices of the prescribed diffusivity fields, predicted speed magnitude, and predicted permeability. Despite pronounced differences in the spatial organization of the diffusivity maps, the predicted speed and permeability fields remain highly consistent across cases. High- and low-valued regions are preserved, and no visible artifacts or spurious structures arise due to diffusivity heterogeneity. Differences between cases are subtle and spatially localized.

Taken together, these results demonstrate that while the model responds to changes in the *mean* diffusivity in a physically consistent manner, it is robust to spatial heterogeneity when the mean value is preserved. This analysis supports the methodological robustness of the framework and justifies the use of a uniform diffusivity in subsequent analyses.

### Predicted results are robust to Gaussian noise but sensitive to sparse outliers

We next evaluated the robustness of the proposed framework to measurement noise in the concentration data. Because DCE-MRI measurements are inherently subject to noise and potential artifacts, it is critical to assess how different noise characteristics propagate into the predictions. Using synthetic concentration fields generated from a 3D transient fluid dynamics simulation, we systematically introduced either distributed Gaussian noise or sparse, high-magnitude outliers, and retrained the model independently for each noise scenario.

Figure 8 summarizes the results. The first column shows clean synthetic concentration data at early (5 min), intermediate (30 min), and late (80 min) times, along with the corresponding predicted permeability. The remaining columns display results obtained after contaminating the concentration data with different types of noise. For Gaussian noise, zero-mean perturbations were added at all spatial locations and time points, with standard deviations equal to 10% or 50% of the mean concentration. To preserve physical plausibility, negative concentrations arising from noise were clipped to zero. For non-Gaussian noise, sparse outliers were introduced by perturbing randomly selected spatial locations (1% or 5% of the domain) with large amplitudes equal to 2× or 5× the maximum concentration, respectively.

**Figure 8:**
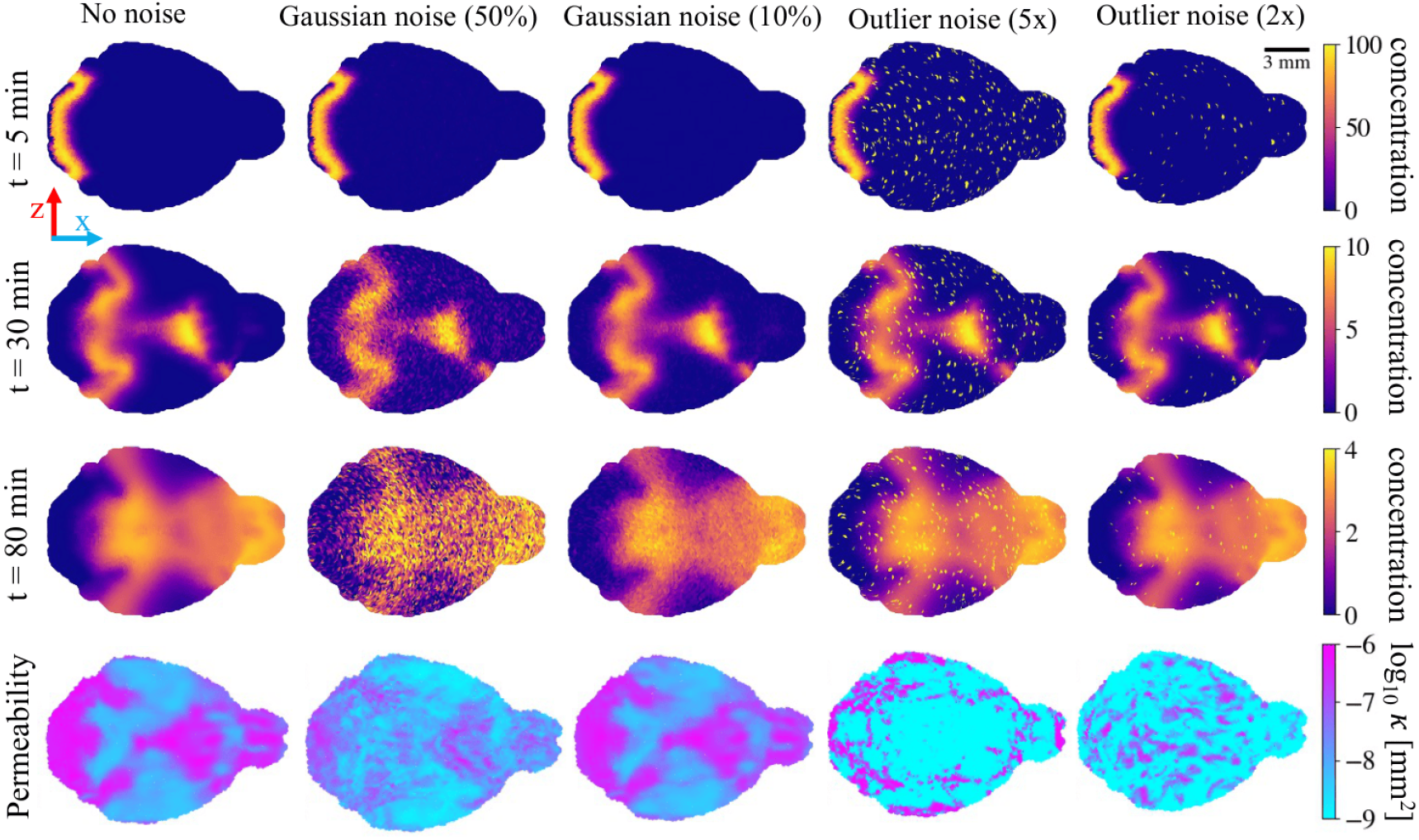
Gaussian noise minimally affects predictions, whereas outliers distort permeability. Synthetic concentration fields are contaminated with Gaussian noise (10% and 50%) and sparse high-magnitude outliers (at 1% and 5% of voxels), and the model is retrained for each case. Rows show concentration snapshots at early, intermediate, and late times, with predicted permeability in the bottom row. Permeability maps are affected little by Gaussian noise, whereas sparse outliers introduce visible distortions. MR-AIV is robust to Gaussian noise and sensitive to outliers.

#### Robustness to distributed Gaussian noise

Gaussian noise has little effect on the predicted permeability fields. For 10% noise, the predicted permeability is nearly indistinguishable from the noise-free case. Even for 50% noise, the model retains the overall spatial structure of permeability, though increased spatial variability and a mild reduction in low-permeability regions are observed. Importantly, no spurious large-scale artifacts emerge, and the dominant permeability patterns are preserved. These results indicate that the framework is robust to distributed, zero-mean Gaussian noise over a broad range of amplitudes.

#### Sensitivity to sparse, high-magnitude outliers

In contrast, localized non-Gaussian noise has a stronger effect. When large-amplitude outliers are introduced at 1% or 5% of spatial locations, noticeable deviations from the noise-free permeability map appear. The effect becomes more pronounced as the outlier magnitude increases (5 ×maximum concentration), leading to visible distortion in permeability patterns. Although the overall range of predicted permeability remains bounded, the spatial organization becomes less consistent with the ground truth, indicating that sparse, high-intensity noise has a stronger impact than Gaussian noise.

#### Identification of failure modes

Comparison reveals a clear distinction between Gaussian noise and outlier noise. The model effectively averages out zero-mean Gaussian perturbations because MR-AIV incorporates a Gaussian negative log-likelihood formulation, which is designed to handle Gaussian-distributed measurement errors under the assumed statistical model. However, outliers violate this Gaussian assumption and introduce localized, non-Gaussian concentration gradients that corrupt the predicted permeability. Importantly, the difference in behaviour arises from the statistical distribution of the noise (Gaussian versus non-Gaussian), rather than simply from its magnitude. These findings highlight a practical limitation: while the framework is robust to Gaussian-distributed measurement noise, it is more vulnerable to uncorrected artifacts or signal spikes. These results support the reliability of the framework with typical imaging noise.

## Conclusion

This study presents a sensitivity analysis and improves the robustness of the MR-AIV framework for inferring brainwide fluid dynamics from DCE-MRI data. By examining the effects of initialization strategies, permeability bounds, signal–concentration modelling assumptions, diffusivity, and measurement noise, we evaluate the stability, physical consistency, and reliability of the method under realistic sources of uncertainty. The results demonstrate robustness across a range of modelling choices. The predicted velocity and permeability fields are affected little by changes to the initial velocity guess, permeability range, SER–concentration mapping, or spatial structure of diffusivity. At the same time, the method exhibits physically consistent sensitivity to the mean diffusivity, confirming that it preserves mechanistic relationships rather than artificially suppressing parameter dependence. Among the tested factors, the initial permeability had the greatest effect on predictions. We introduced a universal, anatomically-informed, ROI-based permeability initialization strategy that significantly improves anatomical alignment and strengthens speed–permeability coherence under Darcy’s law. The framework is robust to moderate levels of distributed Gaussian noise, but not to sparse outliers, underscoring the importance of mitigating outliers during measurement and preprocessing.

Although the speed magnitude was sensitive to the initial permeability guess and mean diffusivity (e.g., the low-speed region is slower in the binary case than in the ROI cases), the spatial locations of the high- and low-speed regions are remarkably similar, as reflected in the high G-SSIM scores. In fact, the G-SSIM score, quantifying the spatial similarity, is above 0.9 (Table 1) for all predicted speed fields across all parameters tested, suggesting that the spatial distribution of the predictions is extremely robust, even though the speed magnitudes are more sensitive.

Brain-wide CSF flow is increasingly implicated in metabolic waste clearance and the pathophysiology of neurodegenerative disease, yet direct measurement of deep-brain flows in vivo has remained out of reach — fuelling fundamental uncertainty about glymphatic function in ageing and neurological disease. MR-AIV addresses this gap by enabling inference of deep-brain flow fields; however, the underlying inverse problem is ill-posed, so the physiological interpretability of its solutions depends critically on prior assumptions. Here, by systematically quantifying the sensitivity of inferred flows to these priors and modelling choices, we establish the conditions and provide concrete best-practice guidelines under which MR-AIV yields robust estimates of the brain-wide fluid velocity field that can be interpreted with confidence. Under these validated conditions, MR-AIV enables direct interrogation of how CSF-mediated clearance varies across brain regions and physiological states, and how these dynamics vary with time-of-day and arousal state and are disrupted in disorders such as Alzheimer’s disease and hypertension. MR-AIV can now directly quantify disease-associated alterations in transport, test mechanistic hypotheses about impaired clearance, and evaluate interventions aimed at restoring normal flow. By converting a previously inaccessible measurement problem into a reproducible and physiologically grounded framework, this work positions MR-AIV as a practical tool for linking brain fluid dynamics to neurological health and therapeutic response.

