## Supplementary material figures for "Robust MR-AIV: A Systematic Study of Robustness Improvement and Sensitivity Analysis of MR-AIV"

A PREPRINT


**Mohammad Vaezi**
Department of Mechanical Engineering
University of Rochester
Rochester, NY 14627, USA

**Juan Diego Toscano**
Division of Applied Mathematics
Brown University
Providence, RI 02912, USA

**Yisen Guo**
Department of Mechanical Engineering
University of Rochester
Rochester, NY 14627, USA

**Ryszard Stefan Gomolka**
Center for Translational Neuromedicine
University of Copenhagen
2200 Copenhagen N, Denmark

**George Em. Karniadakis**
Department of Mechanical Engineering
University of Rochester
Rochester, NY 14627, USA

**Douglas H. Kelley**
Department of Mechanical Engineering
University of Rochester
Rochester, NY 14627, USA

**Kimberly A. S. Boster***
Department of Mechanical Engineering
University of Rochester
Rochester, NY 14627, USA


April 14, 2026

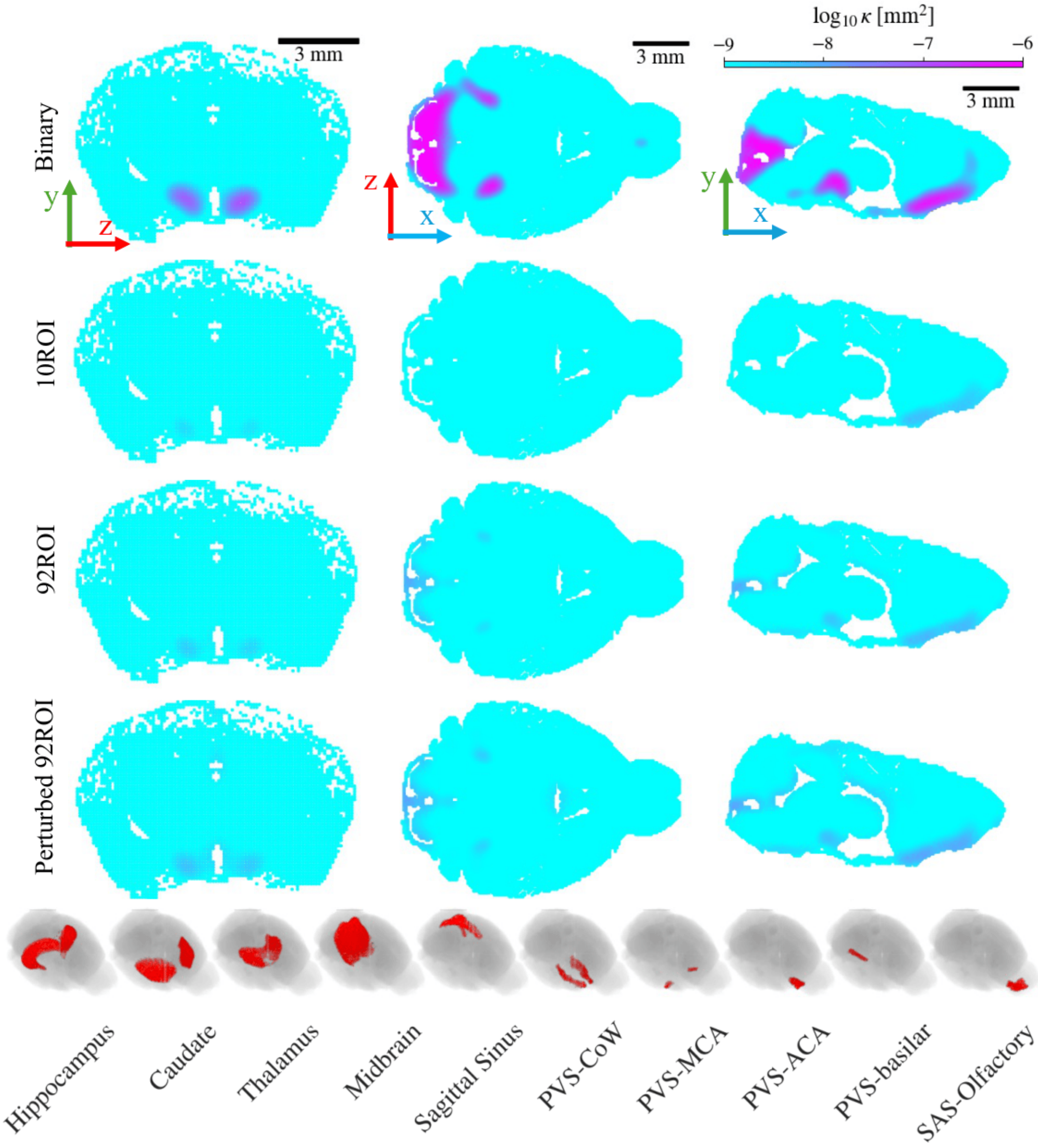


Figure 1: **Initial permeability maps used for model initialization.** Three orthogonal slices (coronal, axial, and midsagittal) of the initial permeability guesses are shown. The binary case is generated based on the early tracer arrival map. The 10ROI and 92ROI maps represent universal initializations derived from trained permeability fields of five wild-type mice. The perturbed 92ROI map is a modified version of the 92ROI initialization used to assess robustness to perturbations. The last row illustrates the 10 regions of interest from the atlas map used to construct the 10ROI initialization, which was generated from permeability fields trained on five wild-type mice that were originally initialized using the early tracer arrival map. Identical initialization of the velocity field is used for all cases.

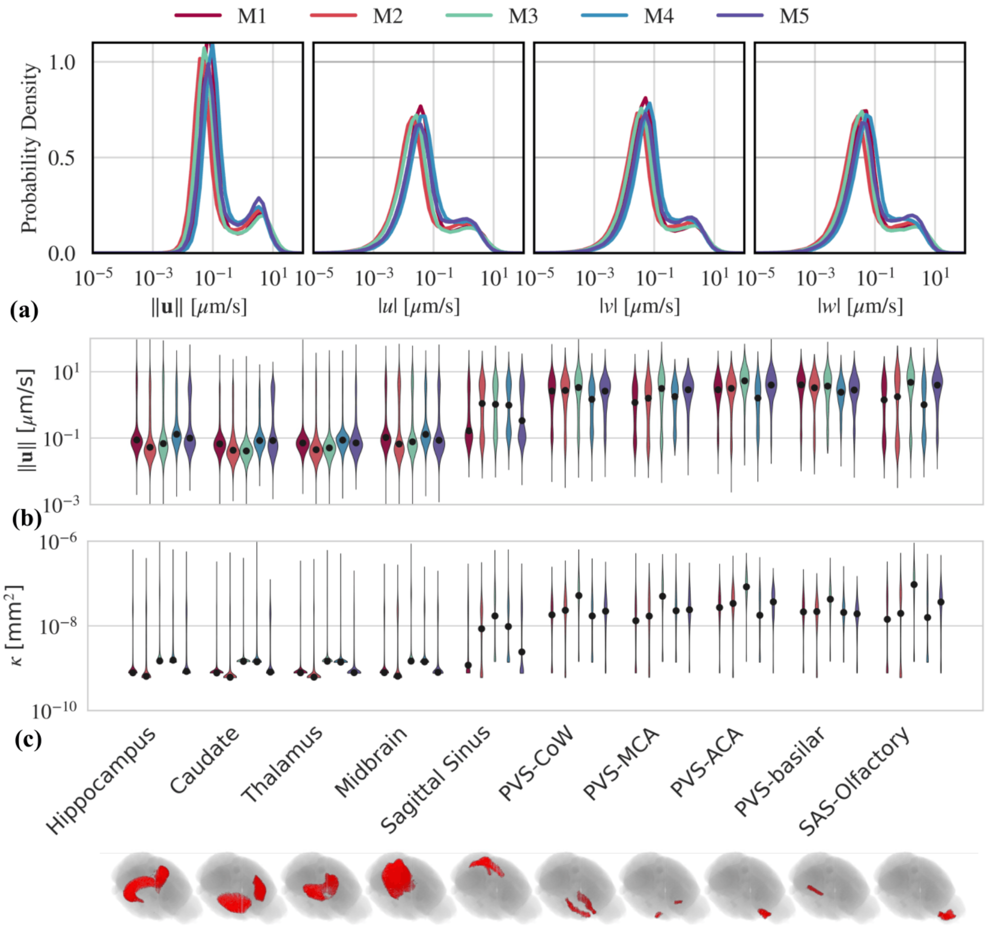


Figure 2: **Speed and permeability based on universal 10ROI permeability map initialization.** We trained permeability based on the early tracers' arrival map for five wild-type mice and mapped them on a standard atlas map to find a universal permeability map. Finally, we used the 10ROI universal map of permeability as initial guess of permeability and trained five different models for five different mice. Panel (a) shows the speed and different components of the velocity for five different wild-type mice. Panels (b) and (c) show speed and permeability distribution at 10 regions of interest (last row of the figure).

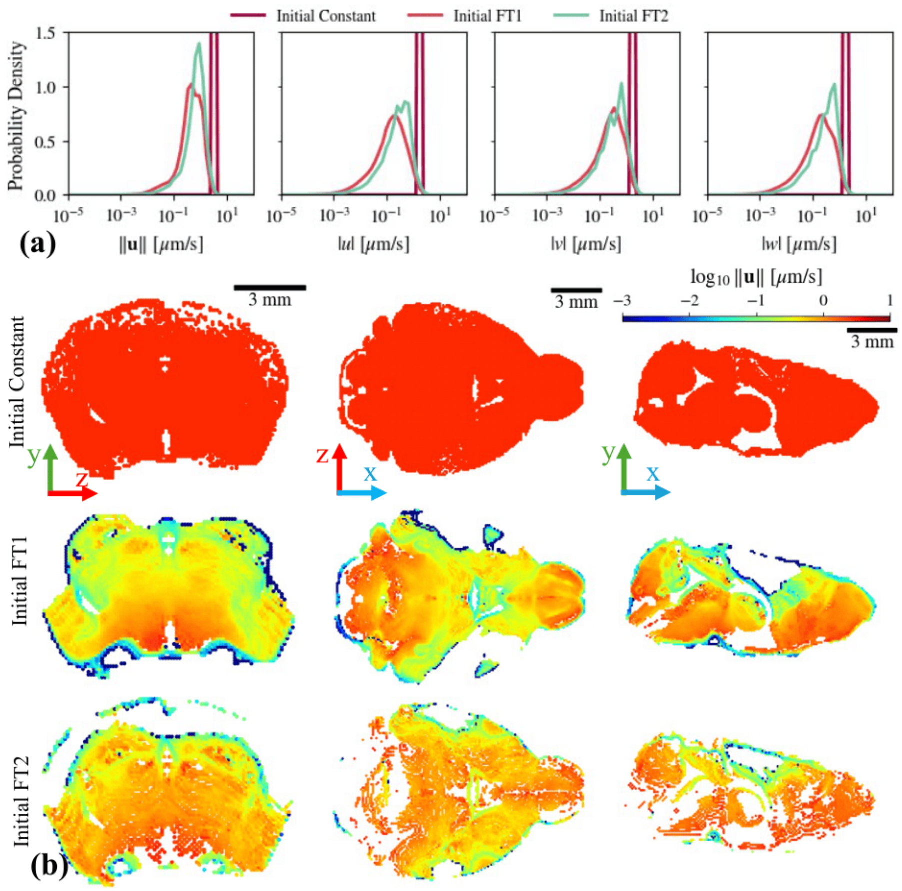


Figure 3: **Comparison of velocity initialization strategies.** Panel (a) compares the distributions of speed and the three Cartesian velocity components ($u$, $v$, and $w$) for three different initialization strategies: a spatially uniform constant-speed field, and two front-tracking–based initializations (FT1 and FT2) obtained using different thresholding criteria. Panel (b) shows three orthogonal slices (coronal, axial, and midsagittal) of the corresponding initial speed maps.

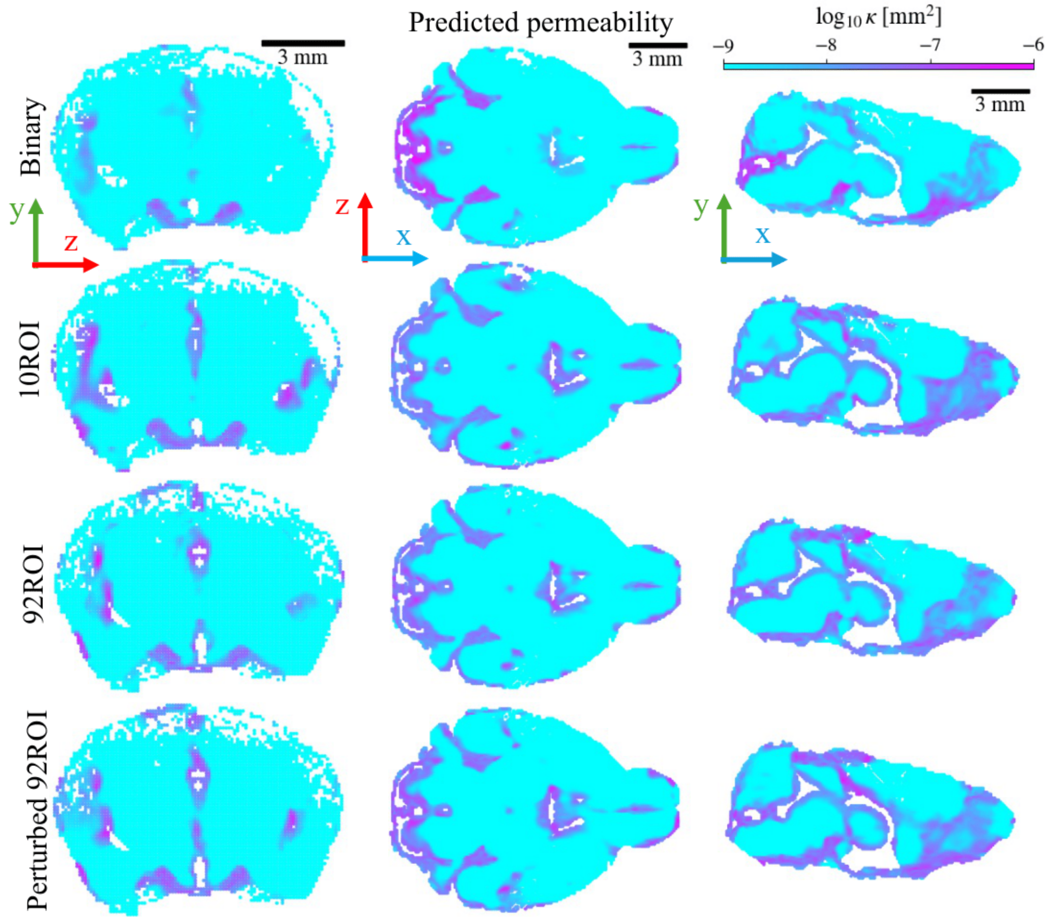


Figure 4: **Predicted permeability fields under different initialization strategies.** Three orthogonal slices (coronal, axial, and midsagittal) of the predicted permeability are shown for models initialized using the binary, 10ROI, 92ROI, and perturbed 92ROI permeability maps. Identical initialization of the velocity field is used for all cases.

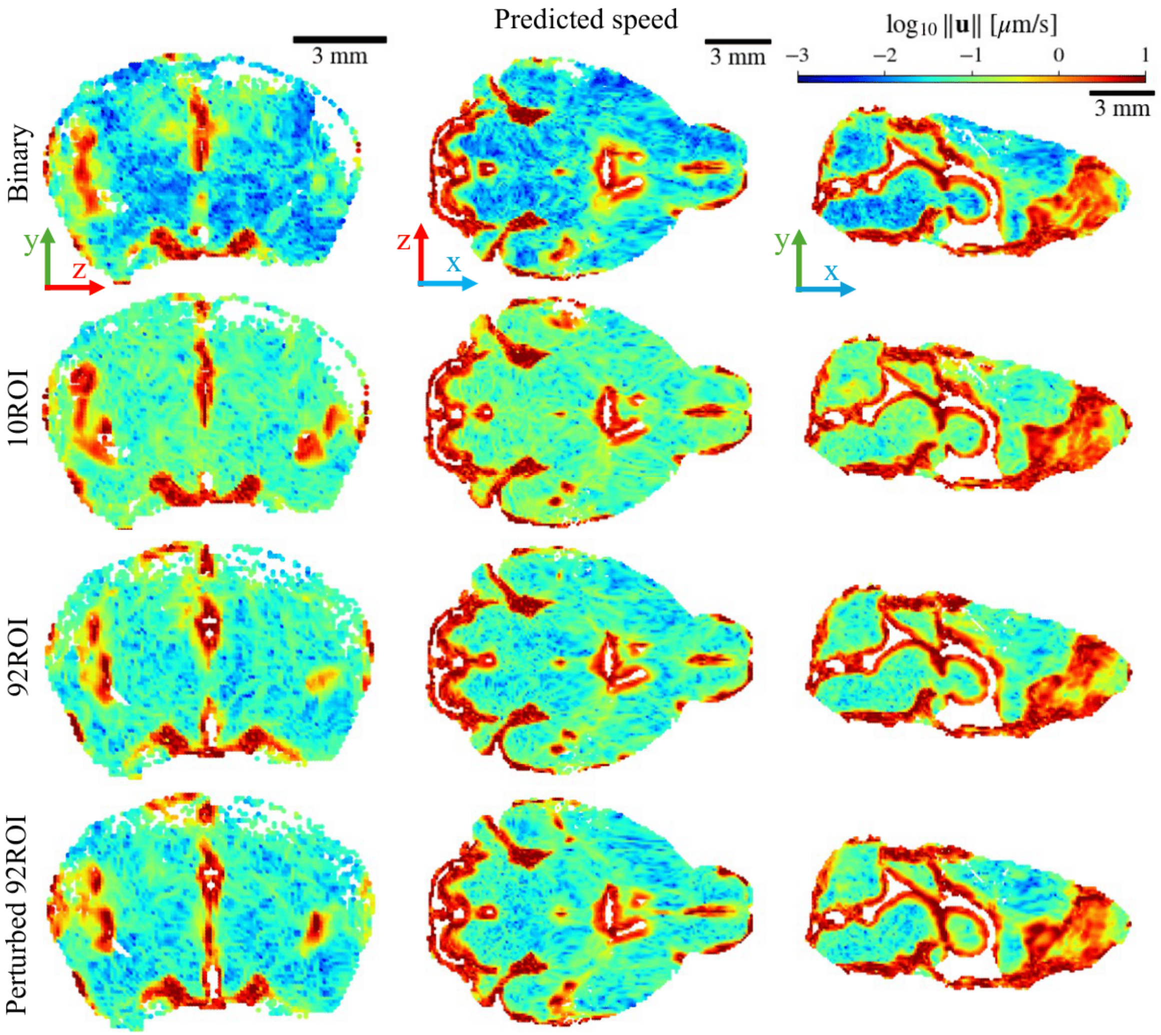


Figure 5: **Predicted speed fields under different permeability initialization strategies.** Three orthogonal slices (coronal, axial, and midsagittal) of the predicted speed are shown for models initialized using the different permeability initialization strategies. Identical initialization of the velocity field is used for all cases.

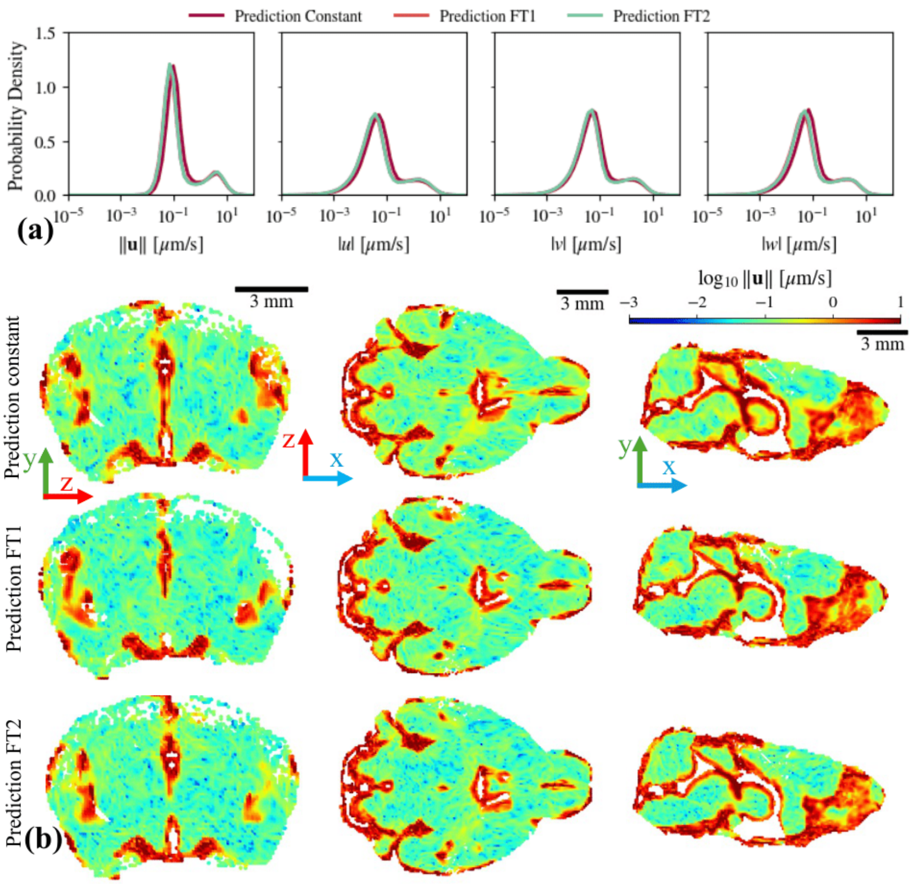


Figure 6: **Predicted velocity fields under different velocity initialization strategies.** Panel (a) shows the distributions of the trained speed and the three Cartesian velocity components ($u$, $v$, and $w$) obtained from three different velocity initialization strategies. Panel (b) presents three orthogonal slices (coronal, axial, and midsagittal) of the corresponding trained speed fields. An identical 10ROI-based permeability initialization is used in all cases.

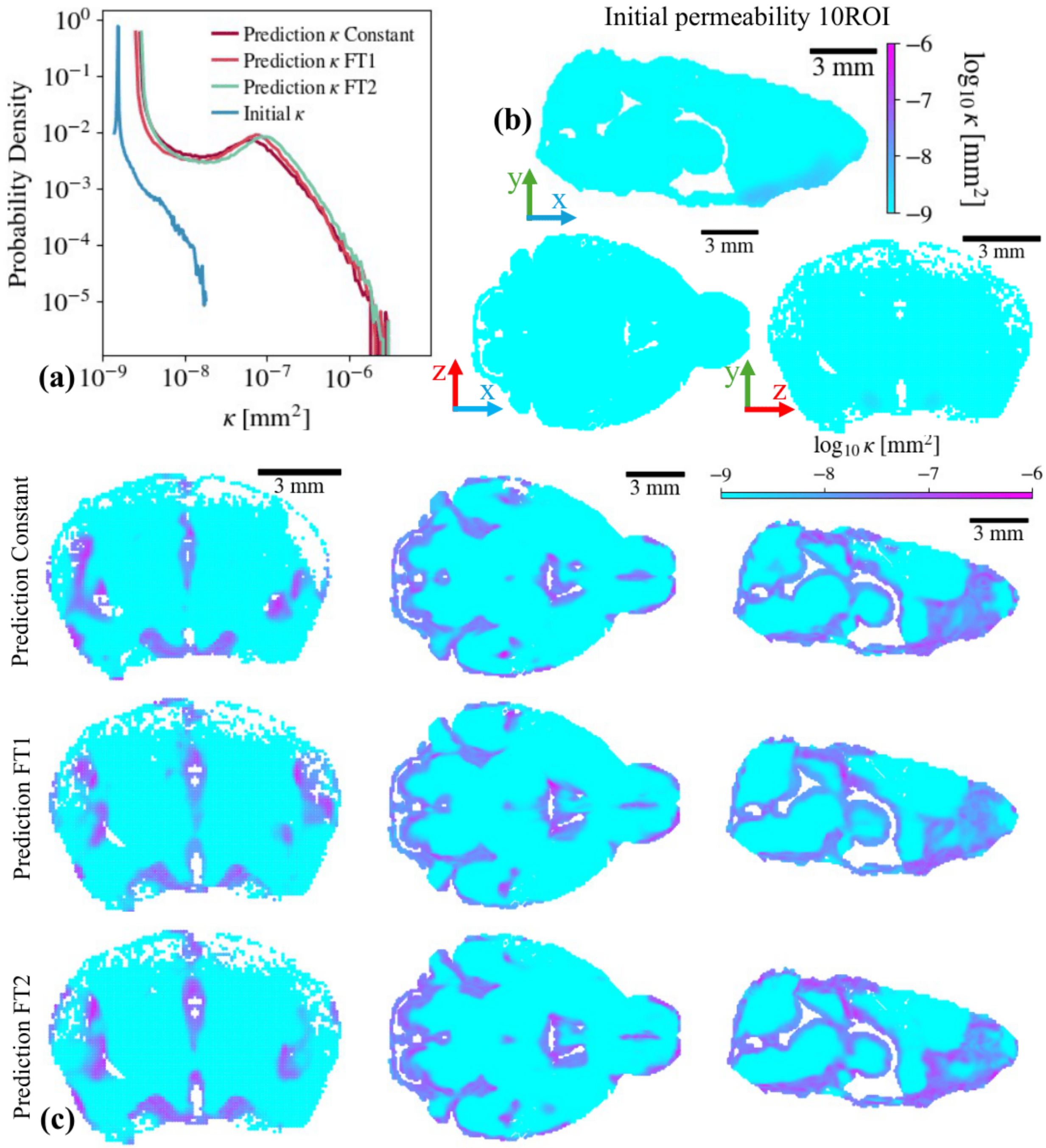


Figure 7: **Predicted permeability under different velocity initialization strategies.** Panel (a) compares the distributions of the predicted permeability obtained from three different velocity initialization strategies, along with the 10ROI initial permeability map used in all cases. Panel (b) shows three orthogonal slices (coronal, axial, and midsagittal) of the 10ROI initial permeability map. Panel (c) presents the corresponding three orthogonal slices of the predicted permeability fields resulting from the three velocity initialization strategies.

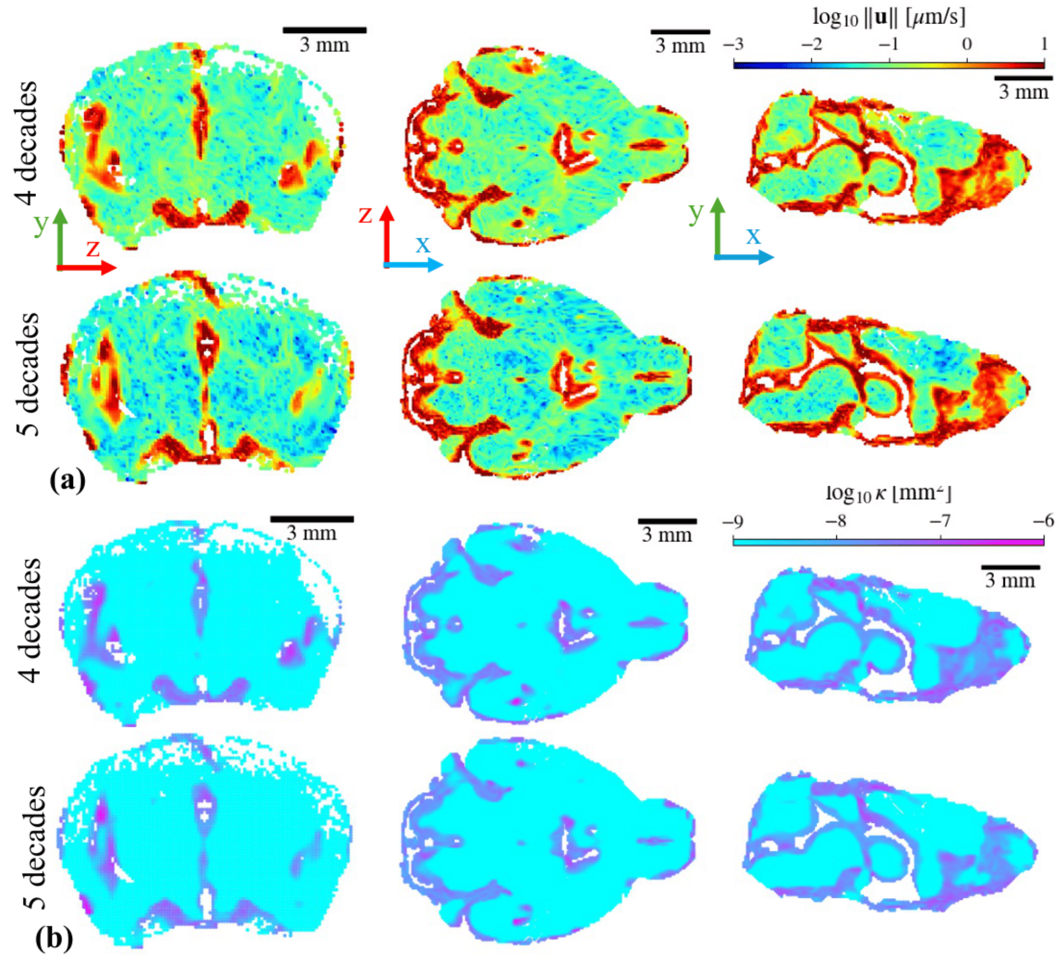


Figure 8: **Effect of permeability learning range on inferred fields.** Panel (a) shows three orthogonal slices (coronal, axial, and midsagittal) of the trained speed obtained when the permeability is learned over ranges spanning four and five orders of magnitude. Panel (b) presents the corresponding three slices of the predicted permeability under the same conditions.

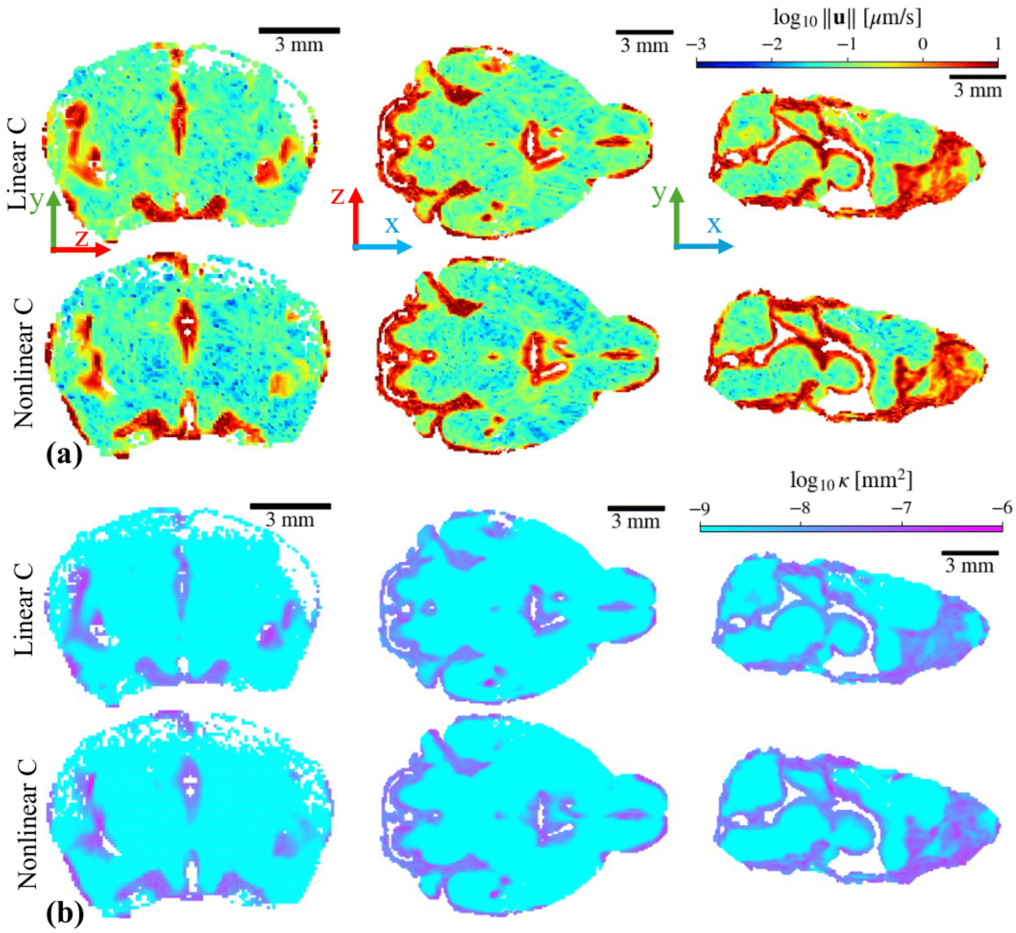


Figure 9: **Effect of SER–concentration modeling on inferred fields.** Panel (a) shows three orthogonal slices (coronal, axial, and midsagittal) of the predicted speed obtained using linear and nonlinear SER–concentration relationships. Panel (b) presents the corresponding three slices of the predicted permeability for the same cases.

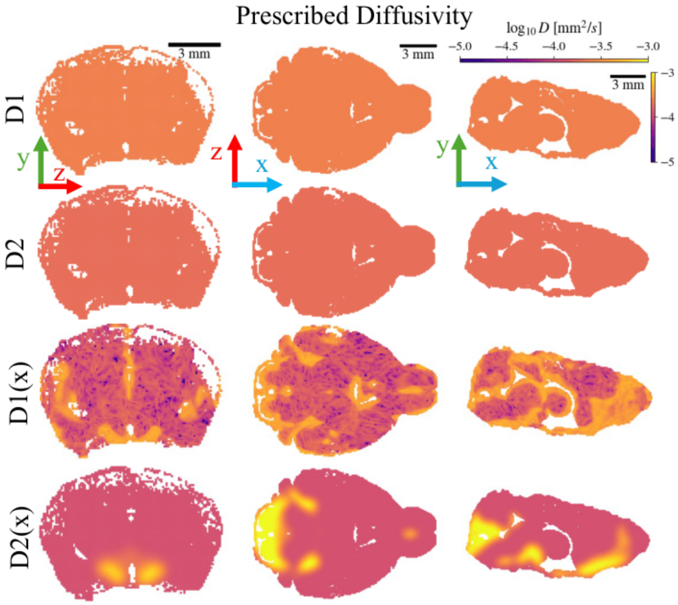


Figure 10: **Prescribed diffusivity fields used in the sensitivity analysis.** This figure shows the four prescribed diffusivity cases considered in the study: two constant diffusivity values ($D_1$ and $D_2$), and two spatially heterogeneous diffusivity fields, $D_1(x)$ and $D_2(x)$, derived from the trained speed map and the early tracer arrival map, respectively.

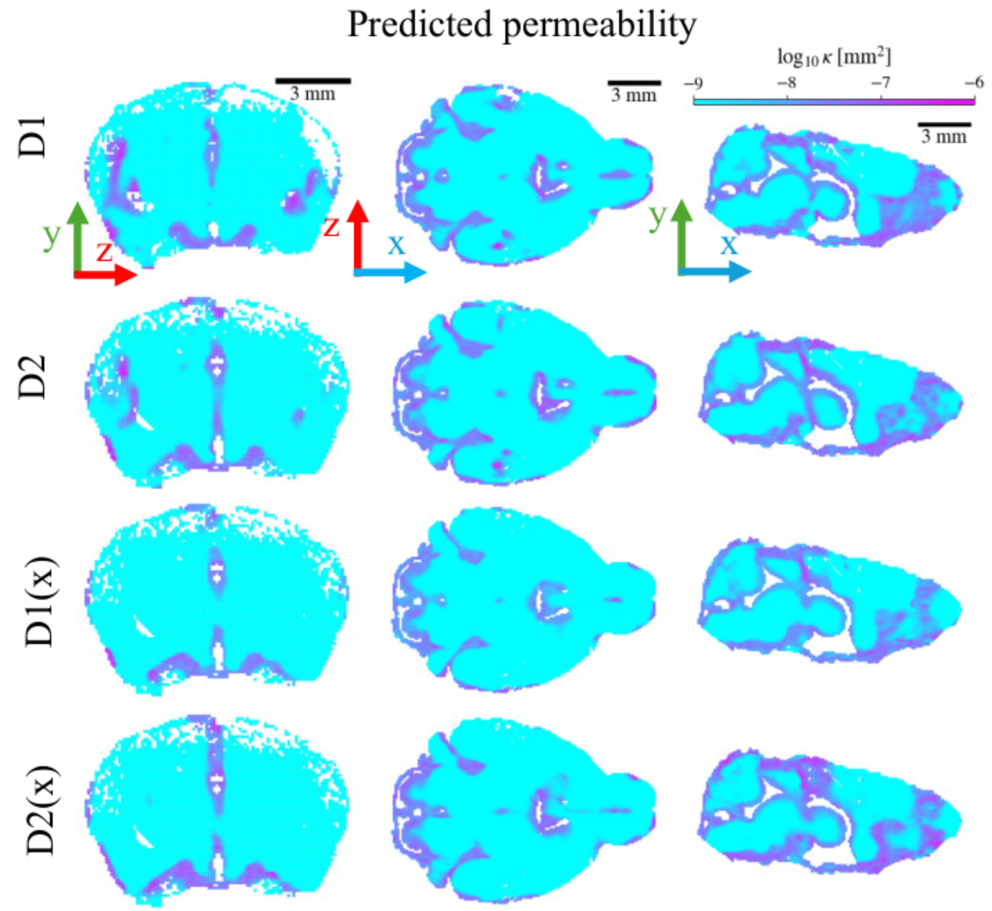


Figure 11: **Predicted permeability under different diffusivity assumptions.** Three orthogonal slices (coronal, axial, and midsagittal) of the trained permeability are shown for four prescribed diffusivity cases: two constant diffusivity values ($D_1$ and $D_2$), and two spatially heterogeneous diffusivity fields ($D_1(x)$ and $D_2(x)$).

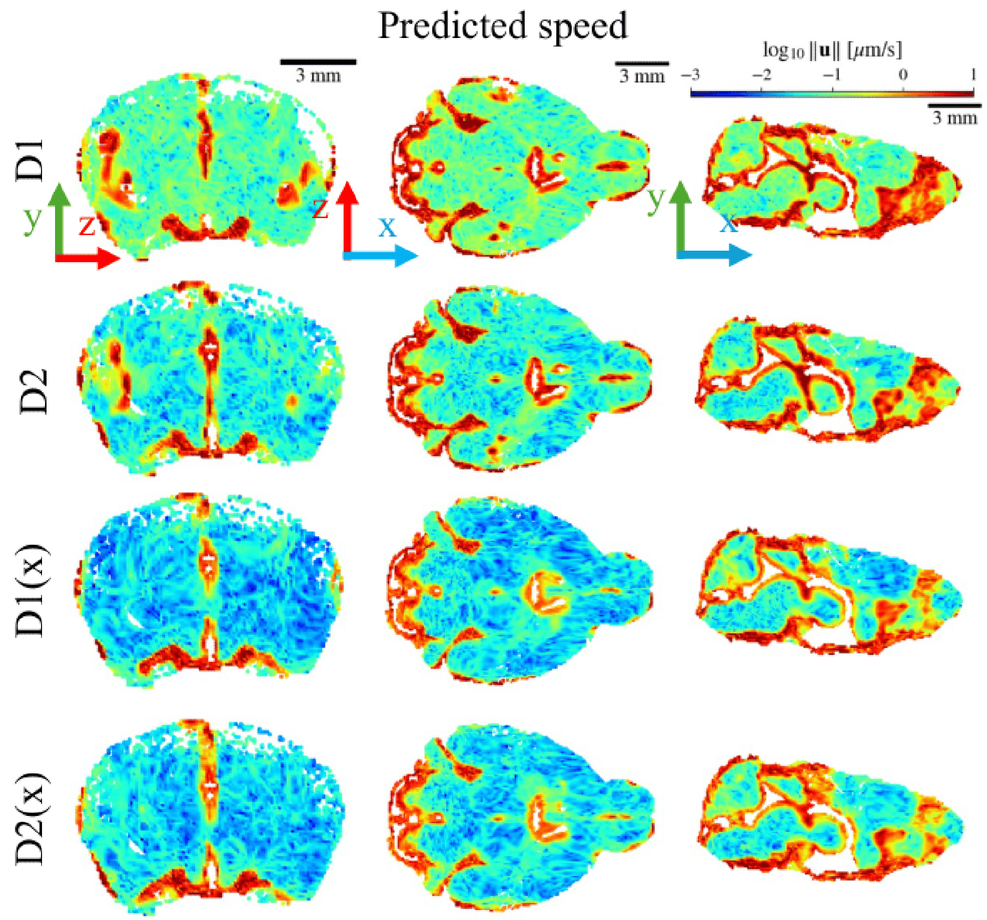


Figure 12: **Predicted speed under different diffusivity assumptions.** Three orthogonal slices (coronal, axial, and midsagittal) of the trained speed are shown for four prescribed diffusivity cases: two constant diffusivity values ($D_1$ and $D_2$), and two spatially heterogeneous diffusivity fields ($D_1(x)$ and $D_2(x)$).
